# Two independent translocation modes drive neural stem cell dissemination into the human fetal cortex

**DOI:** 10.1101/2025.01.08.631865

**Authors:** Ryszard Wimmer, Pauline Lestienne, Clarisse Brunet, Laure Coquand, Amandine Di Cicco, Christophe Chehade, Annasara Artioli, Matthieu Cortes, Anne-Sophie Macé, Xiuyu Wang, Jean-Baptiste Manneville, Bettina Bessière, Ananya Roy, Karin Forsberg-Nilsson, Julia Ladewig, Fabien Guimiot, Alexandre D. Baffet

**Affiliations:** Institut Curie, CNRS UMR144, PSL Research University, Sorbonne University, Paris, France; Department of Translational Brain Research, Central Institute of Mental Health, Medical Faculty Mannheim, Heidelberg University, Mannheim, Germany; Hector Institute for Translational Brain Research (HITBR gGmbH), Mannheim, Germany; German Cancer Research Center (DKFZ), Heidelberg, Germany; UMR 144-Cell and Tissue Imaging Facility (PICT-IBiSA), CNRS-Institut Curie, Paris, France; Laboratoire Matières et Systèmes Complexes, Université Paris Cité, CNRS UMR7057, 10 Rue Alice Domon et Léonie Duquet, F-75013, Paris, France; UF Embryofœtopathologie, Hopital Necker-enfants malades, Paris, France; Department of Immunology, Genetics and Pathology, Uppsala University, Uppsala 751 85, Sweden. Science for Life Laboratory, Uppsala 752 37, Sweden; Science for Life Laboratory, Uppsala 752 37, Sweden; UF de Fœtopathologie – Université de Paris et Inserm UMR1141, Hôpital Robert Debré, Paris, France; Institut national de la santé et de la recherche médicale (Inserm)

## Abstract

The strong size increase of the human neocortex is supported both by the amplification and the basal translocation of a neural stem cell population, the basal radial glial cells (or bRG cells). Using live imaging of second trimester human fetal tissue and cortical organoids, we identify two independent translocation modes for bRG cell colonization of the human neocortex. On top of an actomyosin-dependent movement called mitotic somal translocation (MST), we identify a microtubule-dependent motion occurring during interphase, that we call interphasic somal translocation (IST). We show that IST is driven by the LINC complex, through the nuclear envelope recruitment of the dynein motor and of its activator LIS1. Consequently, IST severely altered in LIS1 patient-derived cortical organoids. We also demonstrate that MST occurs during prometaphase and is a mitotic spindle translocation event. MST is controlled by the mitotic cell rounding molecular pathway, via Moesin and Vimentin, driving translocation. We report that 85% of bRG cell translocation is due to IST, for a total movement of 0,67 mm per month of human fetal gestation. Our work identifies how bRG cells colonize the human fetal cortex, and further shows that IST and MST are conserved in bRG-related migrating glioblastoma cells.

## Introduction

The neocortex is a mammalian-specific brain structure that has expanded dramatically during recent primate evolution. This has culminated in Humans and is associated with the appearance of higher cognitive abilities (Herculano-Houzel, 2012, Fernández and Borrell, 2023). Neocortex expansion is largely supported by the neural stem cells - called the radial glial cells - that generate all neurons, astrocytes, and oligodendrocytes (Allen et al., 2022; Delgado et al., 2022; Del-Valle-Anton et al., 2024; He et al., 2024; Li et al., 2023; Liu et al., 2023; Uzquiano et al., 2022). Two major types of radial glial cells co-exist, with very different abundance and properties across mammals (Fernández et al., 2016) (Figure 1A). Apical radial glial (aRG, also known as vRG) cells are common to all mammals and lie along the lateral ventricles where they form a neuroepithelium (Malatesta et al., 2000; Noctor et al., 2001). At mid-neurogenesis, human aRG cells lose their connection to the pial surface, becoming truncated radial glia (tRGs) (Nowakowski et al., 2016). Basal radial glial cells (bRGs, also known as outer radial glia (oRGs)) on the other hand, are rare and poorly proliferative in certain species such as mice, but extremely abundant and proliferative in others, including primates or ferrets (Fietz et al., 2010; Hansen et al., 2010; Reillo et al., 2011; Sauerland et al., 2018; Matsumoto et al., 2020). They furthermore appear to correlate with the degree of cortical folding (Fernández et al., 2016). bRG cells are particularly numerous in Humans where they are believed to underlie massive neocortical expansion. bRG cells are born from aRG cells and have delaminated, either through mitotic spindle rotation or interphasic loss of adherens junctions (Ostrem et al., 2014; Martínez-Martínez et al., 2016). They form a second stem cell niche, located basally, called the outer subventricular zone (oSVZ) (Liu and Zhang, 2011). How the human oSVZ grows during the course of fetal development to support enhanced neuronal and glial cell production is only starting to be addressed.

**Figure 1.**
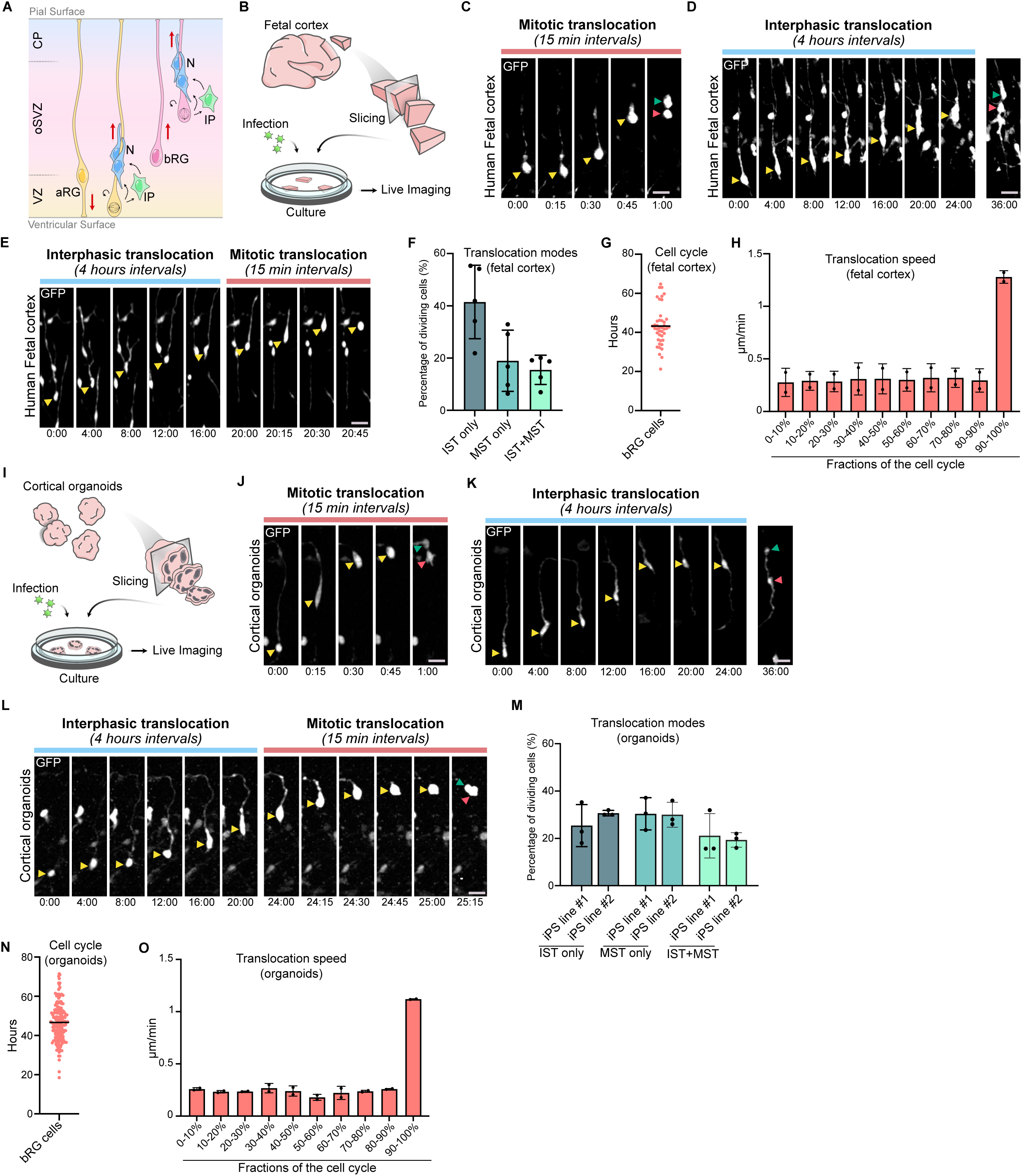
Slow Interphasic and fast mitotic somal translocation. (A) Schematic representation of human neocortex development. VZ: Ventricular Zone. iSVZ and oSVZ: inner and outer Subventricular Zones. CP: Cortical plate. aRG: apical Radial Glial. bRG: basal Radial Glia. IP: Intermediate Progenitor. N: Neuron. (B) Procedure for human fetal cortex slicing, culture, infection and live imaging. (C) Live imaging of a GFP-expressing bRG cell performing mitotic somal translocation in human fetal tissue at pcw 14. (D) Live imaging of a GFP-expressing bRG cell performing interphasic somal translocation in human fetal tissue at pcw 14. (E) Live imaging of a GFP-expressing bRG cell performing interphasic somal translocation followed by mitotic somal translocation in human fetal tissue at pcw 15. (F) Fraction of bRG cells undergoing IST only, MST only or both in human fetal tissue at pcw 12-19 (N=5 fetal samples, 476 bRG cells). (G) Cell cycle duration of bRG cells in human fetal tissue at pcw 16. Cells were live imaged for 92 hours and cell cycle length was measured between two divisions (N=2 fetal samples, 40 cells). (H) Speed of bRG cell somal translocation throughout the cell cycle, between two consecutive divisions, in human fetal tissue pcw 16. The cell cycle was binned into 10 fractions (N=2 fetal samples, 40 cells). (I) Procedure for human cortical organoid slicing, culture, infection and live imaging. (J) Live imaging of a GFP-expressing bRG cell performing mitotic somal translocation in a week 8 human cortical organoid. (K) Live imaging of a GFP-expressing bRG cell performing interphasic somal translocation in a week 8 human cortical organoid. (L) Live imaging of a GFP-expressing bRG cell performing interphasic somal translocation followed by mitotic somal translocation in a week 9 human cortical organoid. (M) Fraction of bRG cells undergoing IST only, MST only or both in week 7-10 human cortical organoids (N=3 organoid batches per iPS line, 314 bRG cells). (N) Cell cycle duration of bRG cells in week 10-11 human cortical organoids. Cells were live imaged for 72 hours and cell cycle length was measured between two divisions (N= 2 organoid batches, 186 bRG cells). (O) Speed of bRG cell somal translocation throughout the cell cycle, between two consecutive divisions, in week 10-11 human cortical organoids. The cell cycle was binned into 10 fractions (N= 2 organoid batches, 186 bRG cells). Yellow arrowheads indicate bRG cell soma, and green and red arrowheads indicate daughter cells. Data are presented as mean values +/- SD. Scale bar = 20 µm. All live imaging montages are in hours:minutes.

Recent evidence has documented the high proliferative potential of human RG cells (Coquand et al., 2024; Lindenhofer et al., 2024). oSVZ growth appears dominated by bRG cell symmetric self-amplifying divisions, coupled to a lower rate of self-consuming neurogenic divisions (Coquand et al., 2024). In cortical organoids, various treatments, such as optimized oxygenation or Leukemia Inhibitory Factor (LIF) stimulation, increase the size of the oSVZ (Andrews et al., 2023; Qian et al., 2020; Walsh et al., 2024). Likewise, overexpression of human-specific genes leads to the expansion of the bRG population and of the oSVZ (Fiddes et al., 2018; Suzuki et al., 2018; Fischer et al., 2022; Pinson et al., 2022; Xing et al., 2024). Importantly, all these amplification events must be coupled to bRG cell dissemination into the growing oSVZ.

Most bRG cells are attached to the pial surface via an elongated process that can be several millimetres long. Once per cell cycle, prior to division, they undergo a so-called mitotic somal translocation (MST), whereby their soma rapidly advances within the process over dozens of micrometres (Hansen et al., 2010) (Figure 1A). This phenomenon was proposed to underly the basal dissemination of bRG cells and the expansion of the oSVZ. bRG cells can adopt various morphologies, with a basal process (the most frequent shape), an apical process, two processes, or no process at all (Betizeau et al., 2013; Kalebic et al., 2019; Coquand et al., 2024). Consequently, MST can occur either basally or apically. MST is furthermore maintained in Pre-Oligodendrocyte progenitor cell (Pre-OPCs), but over shorter distances (Huang et al., 2020).

In aRG cells, the nucleus oscillates along the cell cycle, a process known as Interkinetic Nuclear Migration (INM) (Wimmer and Baffet, 2023). This process is microtubule-dependent, with G1 basal movement - away from the apical centrosome - powered by the kinesin KIF1A, and apical movement - towards the centrosome - powered by dynein and its activator LIS1 (Tsai et al., 2010). In migrating neurons, a nuclear translocation process also occurs, pulled by a leading centrosome (Gonçalves et al., 2020). This movement also relies on the dynein/LIS1 complex as well as on actomyosin contractility (Tsai et al., 2007). However, while in migrating neurons dynein is recruited to the nuclear envelope via the LINC complex in a cell cycle-independent manner, it binds to the nuclear pore complex specifically during G2 in aRG cells (Baffet et al., 2015; Hu et al., 2013).

In bRG cells, MST is dependent on actomyosin contractility and the Rho-associated protein kinase (ROCK), but does not require microtubules (Ostrem et al., 2014), suggesting that it relies on a different mechanism than INM or neuronal migration. mTOR signalling was further shown to modulate MST via a regulation of the actin cytoskeleton and the morphology of bRG cells (Andrews et al., 2020). The exact molecular mechanism of MST, whether the microtubule cytoskeleton also controls other steps of bRG cell translocation, and the contribution of these movements to bRG cell dissemination into the human developing neocortex are however unknown.

Here, using live imaging of human fetal tissue and human cortical organoids, we identify two independent modes of translocation for bRG cells. On top of actomyosin-dependent MST, bRG cells undergo a microtubule-dependent movement during interphase, that we call interphasic somal translocation (IST). IST is slower than MST and is controlled by the LINC complex that recruits the dynein motor and its activator LIS1 to the nuclear envelope for transport. Consequently, IST is affected in LIS1 patient derived organoids. We furthermore show that MST occurs during prometaphase and is therefore a mitotic spindle translocation event. MST is controlled by the mitotic cell rounding pathway, via Ezrin-Radixin-Moesin (ERM) proteins, Vimentin and actomyosin. Both IST and MST are bidirectional and together account for a net bRG cell basal movement of 0,67 mm per month of human fetal gestation. We show that over 85% of this movement is dependent on IST, that is more processive than MST. Finally, we demonstrate that IST and MST are conserved in bRG-related glioblastoma cells and occur through the same molecular pathways. Overall, our work identifies how bRG cells colonize the human fetal neocortex, and how alterations in these mechanisms can be linked to pathological conditions.

## Results

### Fast mitotic and slow interphasic somal translocations

To investigate bRG cell somal translocation, we infected post-conceptional week (pcw) 12-20 human fetal cortical slices with GFP-coding retroviruses and live imaged them for 2 to 4 days (Figure 1B). Typical MST behaviour could be identified in bRG cells, right before cytokinesis (Figure 1C). Remarkably, we could also observe peculiar bRG cell somal translocation events occurring during interphase, albeit at a lower speed (Figure 1D). These cells were *bona fide* bRG cells as they ended up dividing, and live/fixed correlative imaging indicated that cells fixed before division were SOX2+ (Supplementary figure 1A). We named this process interphasic somal translocation (IST), and observed that bRG cells could undergo IST only, MST only, or both (Figure 1E and 1F). We quantified the velocity of somal translocation throughout the entire cell cycle of bRG cells. Live imaging of fetal tissue for 4 days revealed an average cell cycle duration of 43 hours, around two times higher than that of mouse aRG cells (Pilaz et al., 2009) (Figure 1G and Supplementary Figure 1B). Translocation velocity remained constant throughout interphase and greatly accelerated around mitosis, confirming the bi-phasic migration behaviour (Figure 1H). We next performed the same analysis in human week 7-12 cortical organoids (Qian et al., 2016; Sloan et al., 2018), using two independent iPS cell lines (Figure 1I). Interphasic and mitotic translocation events were also observed, occurring at similar frequencies than in fetal tissues (Figure 1J-M). 4-day live imaging revealed a highly similar cell cycle duration for bRG cells (47 hours), as well as comparable velocities, both in interphase and mitosis (Figure 1 N & O). Therefore, human bRG cells translocate slowly throughout interphase, and rapidly during mitosis, a process conserved between human cortical organoids and fetal tissue.

### IST is microtubule dependant and MST is actomyosin dependent

We next asked whether interphasic and mitotic translocations were driven through distinct molecular mechanisms or were merely the same process occurring at different speeds. To test this, we first established *in vitro* 2D cultures of fetal RG cells, using fibronectin-coated dishes (Figure 2A). The cells maintained an elongated morphology, were positive for the RG markers SOX2, HOPX, PTPRZ1 and LIFR and negative for EOMES and NEUN (Supplementary Figure 2A). These cells still performed IST and MST *in vitro*, indicating that these movements are intrinsic features of bRG cells (Figure 2 B-D). Live imaging of these cells between two divisions revealed a ∼2 times shorter cell cycle than in tissues (Figure 2E). IST velocity was constant during interphase, while MST was much faster, as they were in tissues (Figure 2F). However, both IST and MST were substantially faster in 2D dissociated cultures (∼2X), as compared to organoids or fetal tissue, suggesting a negative effect of tissue environment on translocation velocity.

**Figure 2.**
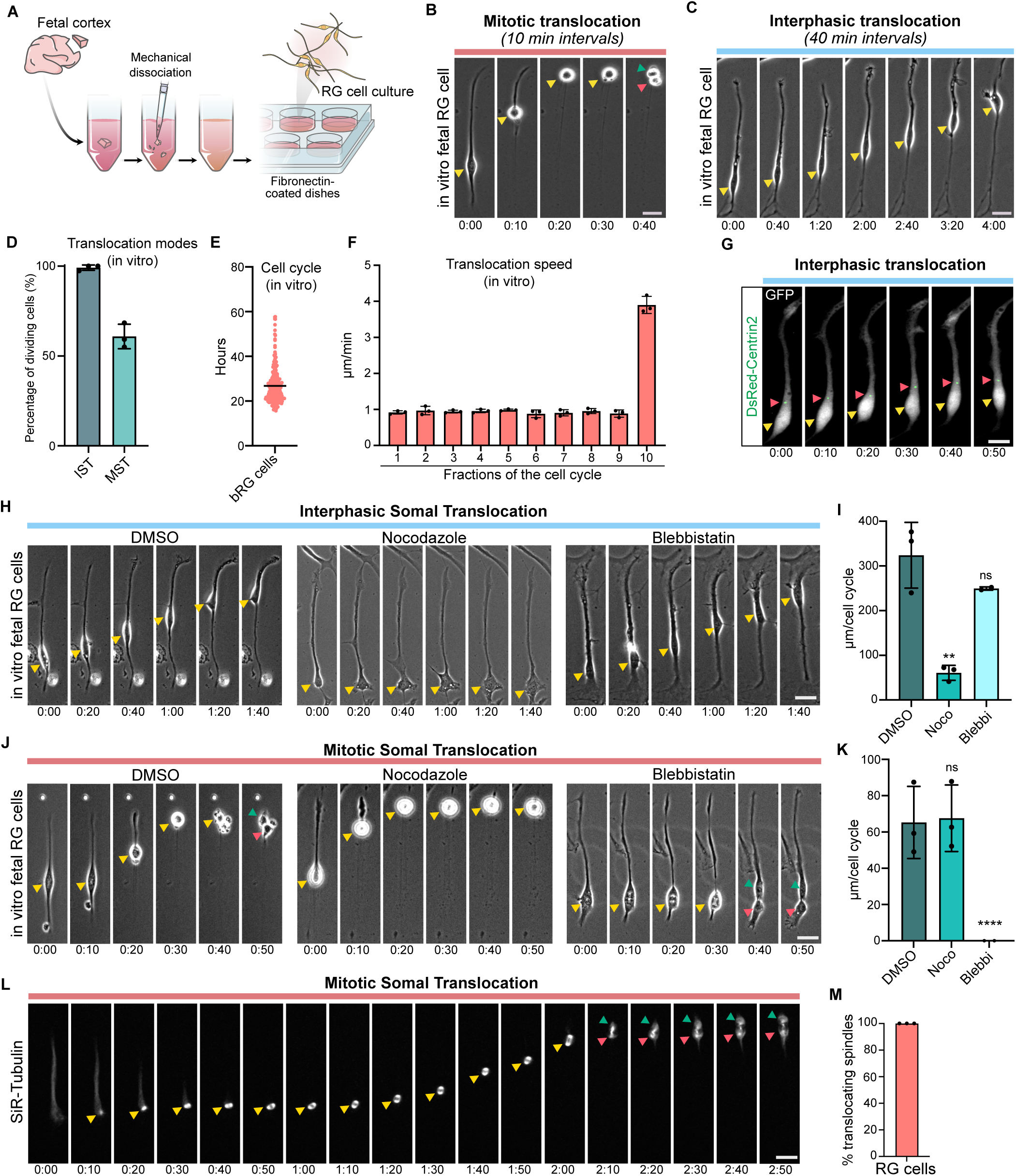
IST is microtubule dependant and MST is actomyosin dependent. (A) Procedure for human fetal cortex dissociation, primary bRG cell 2-dimensional *in vitro* culture (*in vitro* bRG), and live imaging. (B) Live imaging of an *in vitro* bRG cell performing IST. (C) Live imaging of an *in vitro* bRG cell performing MST. (D) Fraction of *in vitro* bRG cells undergoing IST, MST (N=3 experiments, 240 bRG cells). (E) Cell cycle duration of *in vitro* bRG cells. Cells were live imaged for 72 hours and cell cycle length was measured between two divisions (N=3 experiments, 240 bRG cells). (F) Speed of *in vitro* bRG cell somal translocation thorough the cell cycle, between two consecutive divisions. The cell cycle was binned into 10 fractions (N=3 experiments, 240 bRG cells). (G) Live imaging of an *in vitro* bRG cell performing IST and expressing GFP and DsRed-Centrin2. (H) Live imaging of *in vitro* bRG cells during interphase, treated with DMSO, nocodazole (1*μ*M) or blebbistatin (10*μ*M). (I) Quantification of IST amplitude following treatment with DMSO, nocodazole (1*μ*M) or blebbistatin (10*μ*M) in *in vitro* bRG cells (N=3 experiments, 724 bRG cells). (J) Live imaging of *in vitro* bRG cells during mitosis, treated with DMSO, nocodazole (1*μ*M) or blebbistatin (10*μ*M). (K) Quantification of MST amplitude following treatment with DMSO, nocodazole (1*μ*M) or blebbistatin (10*μ*M) in *in vitro* bRG cells (N=3 experiments, 724 bRG cells). (L) Live imaging of an *in vitro* bRG cell RG cell performing MST and expressing SiR-Tublin. Mitotic spindle begins to form before translocation. (M) Quantification of the fraction bRG cells in which mitotic spindle forms before somal translocation starts (N=2 experiments, 94 bRG cells). Yellow arrowheads indicate bRG cell soma, and green and red arrowheads indicate daughter cells. Data are presented as mean values +/− SD. Scale bar = 20 µm. All live imaging montages are in hours:minutes. *p<0,05; **p<0,01; ****p<0,0001, ns: non-significant by two-tailed unpaired t-tests.

To investigate the role of the cytoskeleton in IST and MST, we first expressed the centrosomal marker Centrin2 in dissociated RG cells. During IST, the centrosome always moved ahead of the nucleus and organized a microtubule cage around the nucleus, consistent with microtubule-dependent pulling forces on the nuclear envelope (Figure 2G and supplementary Figure 2B). To test this, as well as for a role of actomyosin contractility, cells were treated with the microtubule depolymerizing drug nocodazole and the myosin II inhibitor blebbistatin. These experiments revealed that IST was severely affected by microtubule depolymerization, but not by actomyosin inhibition (Figure 2H & I). Conversely, and as previously published (Ostrem et al., 2014), MST was abolished by myosin II inhibition but not by microtubule depolymerization (Figure 2J & K). Finally, we asked whether MST is a nuclear translocation event, as IST, or if this movement occurs after nuclear envelope breakdown. To test this, we incubated dissociated RG cells with Sir-Tubulin and performed live imaging. These experiments revealed that the mitotic spindle always formed before MST initiation, indicating that MST is not a nuclear translocation event but a mitotic spindle translocation event, occurring during prometaphase-metaphase (Figure 2L & M). Overall, these results show that IST is a microtubule-dependent nuclear transport event occurring throughout interphase, while MST is an actomyosin-dependent mitotic spindle translocation event occurring during prometaphase.

### Dynein, LIS1, and the LINC complex drive IST in bRG cells

We next addressed the molecular mechanism of IST. We first confirmed that, as in dissociated cultures, nocodazole but not blebbistatin impaired IST in cortical organoids (Supplementary Figure 3A). Because microtubule-dependent nuclear transport is largely dependent on the dynein motor in many cell types, we infected week 8-11 human cortical organoids with dynein heavy chain (DYNC1H1) shRNA-coding lentiviruses. Live imaging revealed that dynein loss of function affected the amplitude of IST, with a 3-fold decrease of the distance travelled over time (Figure 3A & C). These results were obtained for two independent and validated shRNA sequences. The role of dynein for IST was confirmed in human in fetal tissue, in three independent live imaged samples (pcw 16-20) (Figure 3B & E). Dynein loss of function did not affect MST (Figure 3D & F and Supplementary Figure 3B & C). We then knocked down the dynein activator LIS1 in bRG cells. Infection with retroviruses encoding LIS1 shRNA constructs strongly affected IST in live imaged human cortical organoids, fetal tissue, and dissociated fetal RG cultures (Figure 3A-C & E and Supplementary Figures 3D & E). As for dynein, LIS1 knockdown did not affect MST (Figure 3D & F and Supplementary Figures 3B & C). Mutations in the LIS1 gene are the most prevalent cause of lissencephaly in Humans, a disease associated with neuronal positioning defects (Reiner et al., 1993; Iefremova et al., 2017). To test if bRG cell translocation defects may occur in lissencephalic tissue, we generated LIS1 patient-derived cortical organoids, from two affected individuals (Zillich et al., 2022). Live imaging of bRG cells within these organoids at weeks 8-11 confirmed a strong IST alteration, without any MST defect (Figure 3G-I). Therefore, dynein and its activator LIS1 control nucleokinesis during IST in bRG cells.

**Figure 3.**
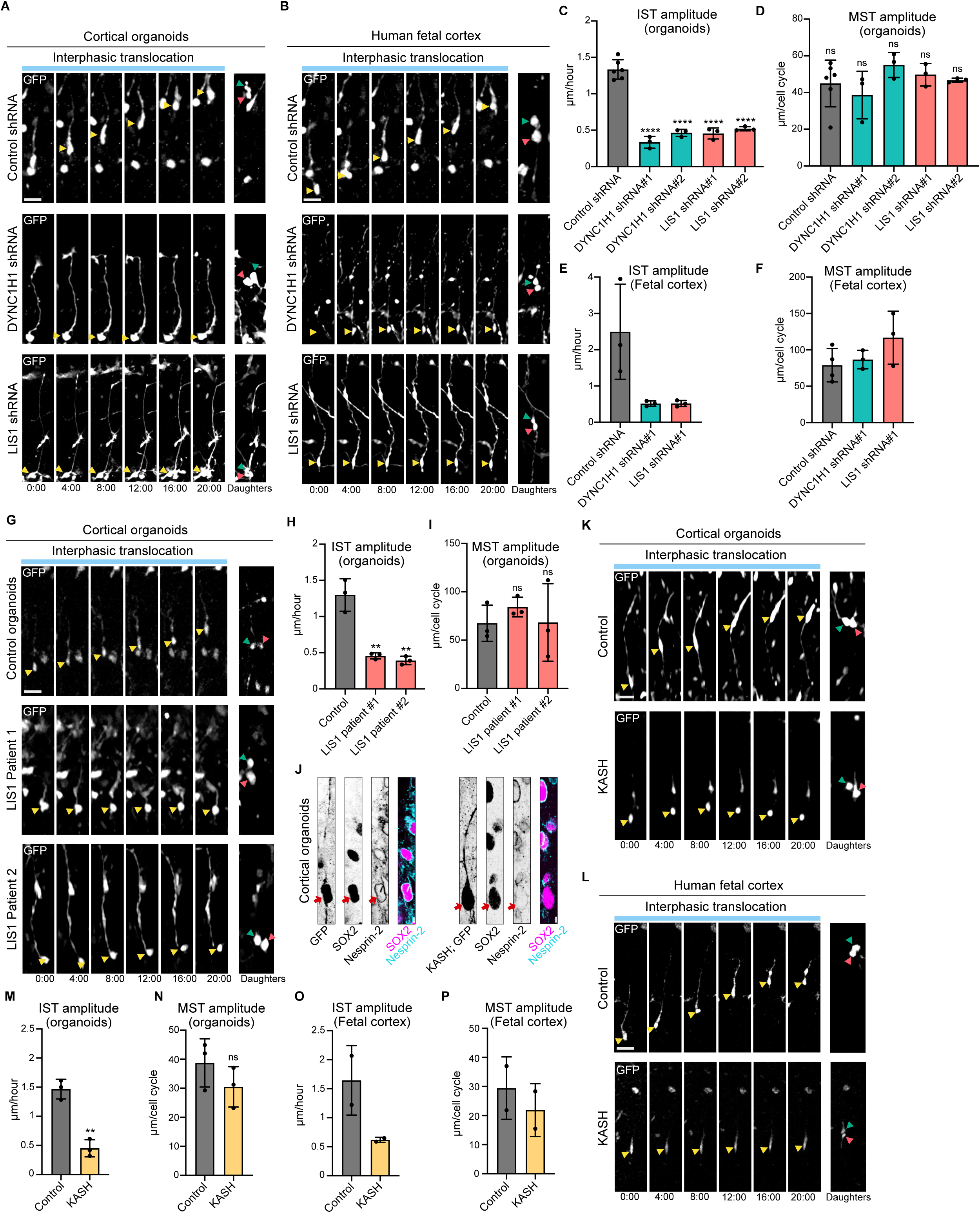
Dynein-LIS1 recruitment to the nuclear envelope by the LINC complex drives IST. (A) Live imaging of interphasic human bRG cells expressing control, DYNC1H1 or LIS1 shRNA constructs in human cortical organoids (week 8-11). shRNA plasmids co-express GFP. (B) Live imaging of interphasic human bRG cells expressing control, DYNC1H1 or LIS1 shRNA constructs in human fetal tissue (pcw 16-18). (C) Quantification of IST amplitude in human bRG cells expressing control, DYNC1H1 or LIS1 shRNA constructs in human cortical organoids (N=3 organoid batches, 899 bRG cells, week 8-11). Two independent shRNA plasmids were used for each knockdown. (D) Quantification of MST amplitude in human bRG cells expressing control, DYNC1H1 or LIS1 shRNA constructs in human cortical organoids (N=3 organoid batches, 899 bRG cells, week 8-11). (E) Quantification of IST amplitude in human bRG cells expressing control, DYNC1H1 or LIS1 shRNA constructs in human fetal tissue (N=3 fetal samples, 385 bRG cells, pcw 16-20). (F) Quantification of MST amplitude in human bRG cells expressing control, DYNC1H1 or LIS1 shRNA constructs in human fetal tissue (N=3 fetal samples, 385 bRG cells, pcw 16-20). (G) Live imaging of interphasic human bRG cells expressing GFP in control cortical organoids and two different patient-derived LIS1-mutated organoids (week 8-11). (H) Quantification of IST amplitude in control cortical organoids and two different patient-derived LIS1-mutated organoids (N=3 organoid batches, 397 bRG cells, week 8-11). (I) Quantification of MST amplitude in control cortical organoids and two different patient-derived LIS1-mutated organoids (N=3 organoid batches, 397 bRG cells, week 8-11). (J) Immunostaining for SOX2 and Nesprin-2 in cortical organoids expressing GFP or the KASH dominant negative together with GFP (week 9). Red arrows indicate nuclear envelope of construct-expressing cells. (K) Live imaging of interphasic human bRG cells expressing control or KASH constructs in human cortical organoids (week 8). KASH plasmid co-expresses GFP. (L) Live imaging of interphasic human bRG cells expressing control or KASH constructs in human fetal tissue (pcw 16). KASH plasmid co-expresses GFP. (M) Quantification of IST amplitude in human bRG cells expressing control or KASH constructs in human cortical organoids (N=3 organoid batches, weeks 8-11, 201 bRG cells). (N) Quantification of MST amplitude in human bRG cells expressing control or KASH constructs in human cortical organoids (N=3 organoid batches, weeks 8-11, 201 bRG cells). (O) Quantification of IST amplitude in human bRG cells expressing control or KASH constructs in human fetal tissue (N=2 fetal samples, pcw 16-18, 40 bRG cells). (P) Quantification of MST amplitude in human bRG cells expressing control or KASH constructs in human fetal tissue (N=2 fetal samples, pcw 16-18, 40 bRG cells). Yellow arrowheads indicate bRG cell soma, and green and red arrowheads indicate daughter cells. Data are presented as mean values +/− SD. Scale bar = 20 µm. All live imaging montages are in hours:minutes. **p<0,01; ****p<0,0001, ns: non-significant by two-tailed unpaired t-tests.

We then asked how the dynein-LIS1 motor complex was targeted to the nuclear envelope of bRG cells. Because IST occurred with a leading centrosome and throughout interphase, we hypothesized that it worked through a similar mechanism than the LINC complex-dependent nuclear migration in neurons, but different from the nuclear pore-dependent INM in aRG cells (Hu et al., 2013). To test this, we expressed the KASH domain of Nesprin-2, which acts as a dominant negative by displacing all Nesprins from the nuclear envelope (Luxton et al., 2010). This construct efficiently displaced Nesprin-2 from the nuclear envelope of bRG cells in cortical organoids (Figures 3J). Live imaging revealed that KASH expression severely altered IST in human cortical organoids, fetal tissue, and fetal RG *in vitro* cultures, without any effect on MST (Figures 3K-P and Supplementary Figure3F). For all these conditions, we noted that IST was affected but not abolished. We therefore wondered whether acto-myosin contractility could be taking over to support partial movement. To test this, we incubated KASH-expressing *in vitro* bRG cells with blebbistatin. This however did not increase the IST phenotype, further indicating that IST is actomyosin-independent (Supplementary figure 3F). Overall, these experiments show that IST is driven by dynein-LIS1 pulling forces, which are recruited to the nuclear envelope by the LINC complex (Figure 4J). This process is molecularly similar to nucleokinesis in migrating neurons, but different from INM in aRG cells.

**Figure 4.**
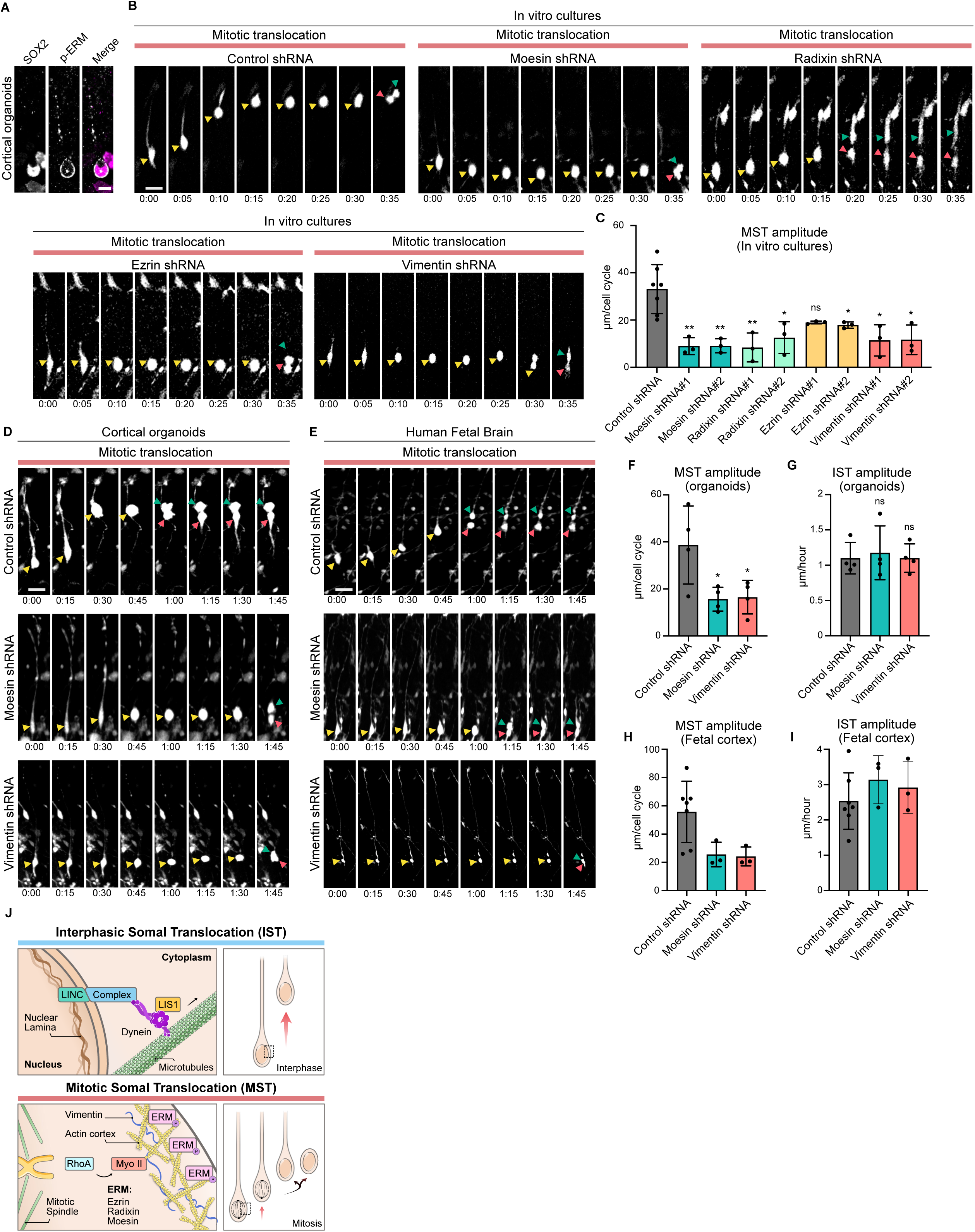
Mitotic Cell Rounding drives Mitotic Somal Translocation in bRG cells. (A) Immunostaining for SOX2 and p-ERM in mitotic bRG cell from week 8 cortical organoid. (B) Live imaging of *in vitro* mitotic human bRG cells expressing control, Moesin, Radixin, Ezrin or Vimentin shRNA constructs. (C) Quantification of IST amplitude in *in vitro* interphasic human bRG cells expressing control, Moesin, Radixin, Ezrin or Vimentin shRNA constructs. Two independent shRNA plasmids were used for each knockdown (N=7 control samples and N=3 sample per shRNA condition, 522 bRG cells). (D) Live imaging of mitotic human bRG cells expressing control, Moesin or Vimentin shRNA constructs in human cortical organoids (week 9). (E) Live imaging of mitotic human bRG cells expressing control, Moesin or Vimentin shRNA constructs in human fetal tissue (pcw 20). (F) Quantification of MST amplitude in human bRG cells expressing control, Moesin or Vimentin shRNA constructs in human cortical organoids (N=3 organoid batches, 453 bRG cells, weeks 8-12). (G) Quantification of IST amplitude in human bRG cells expressing control, Moesin or Vimentin shRNA constructs in human cortical organoids (N=3 organoid batches, 453 bRG cells, weeks 8-12). (H) Quantification of MST amplitude in human bRG cells expressing expressing control, Moesin or Vimentin shRNA constructs in human fetal tissue (N=3 fetal samples, 529 bRG cells, pcw 14-20). (I) Quantification of IST amplitude in human bRG cells expressing expressing control, Moesin or Vimentin shRNA constructs in human fetal tissue N=3 fetal samples, 529 bRG cells, pcw 14-20). (J) Schematic representation of the molecular mechanisms driving IST and MST. Yellow arrowheads indicate bRG cell soma, and green and red arrowheads indicate daughter cells. Data are presented as mean values +/− SD. Scale bar = 20 µm. All live imaging montages are in hours:minutes. *p<0,05; **p<0,01, ns: non-significant by two-tailed unpaired t-tests.

### Mitotic cell rounding drives mitotic Somal translocation in bRG cells

Most cells round up for mitosis to enable proper chromosome segregation, through a process known as mitotic cell rounding (Taubenberger et al., 2020). This process occurs via mitotic phosphorylation of the RhoGEF Ect2 by Cdk1, which activates RhoA to trigger actomyosin contractility, and by mitotic phosphorylation of the ezrin-radixin-moesin (ERM) proteins, that crosslink the actin cortex to the plasma membrane (Kunda et al., 2008; Ramanathan et al., 2015; Toyoda et al., 2017). A vimentin layer beneath the actin cortex further increases mitotic cortical tension (Figure 4J) (Duarte et al., 2019; Serres et al., 2020). We hypothesized here that this cortical tension is then dissipated into the bRG cell elongated process, leading to a translocation of the soma. ERM phosphorylation was indeed detected at the cortex of mitotic bRG cells (Figure 4A). To test for the role of mitotic cell rounding in MST, we first knocked down one by one the members of the ERM family as well as Vimentin in bRG cell *in vitro* cultures using shRNA constructs. Knockdown of Moesin and Radixin strongly reduced the average amplitude of MST, without affecting IST, while knockdown of Ezrin had a milder effect (Figures 4B & C). Knockdown of Vimentin phenocopied Moesin and Radixin loss of function, impairing proper MST in these cells (Figures 4B & C).

We next asked if the roles of Moesin and Vimentin were conserved in 3D samples. Knockdown of Moesin and Vimentin in week 8-12 cortical organoids severely altered MST without affecting IST (Figures 4D, F & G). Finally, we live imaged multiple independent human fetal samples for 48h following control or Moesin and Vimentin shRNA-mediated knockdowns. As in organoids, the loss of function of these mitotic cell rounding factors severely disrupted MST, without affecting IST (Figures 4E, H & I) Together, these results identify mitotic cell rounding as the main driver of MST in human bRG cells, likely via a tension dissipation mechanism into their elongated processes (Figure 4J).

### IST contributes more than MST to bRG cells colonization of the human fetal neocortex

Next, we measured the relative contribution of interphasic and mitotic translocations to bRG cell colonization of the human developing neocortex. The average amplitude of IST and MST movement in fetal tissue, throughout an entire cell cycle and including the static cells, was 77 and 20.4 *μ*M respectively, indicating that IST leads to 3.7x more movement of the bRG cell soma than MST (Figure 5A). A similar bias for IST was observed in cortical organoids, albeit smaller (1.7x) because MST was higher and IST slightly lower than in fetal tissues (Figure 5A).

**Figure 5.**
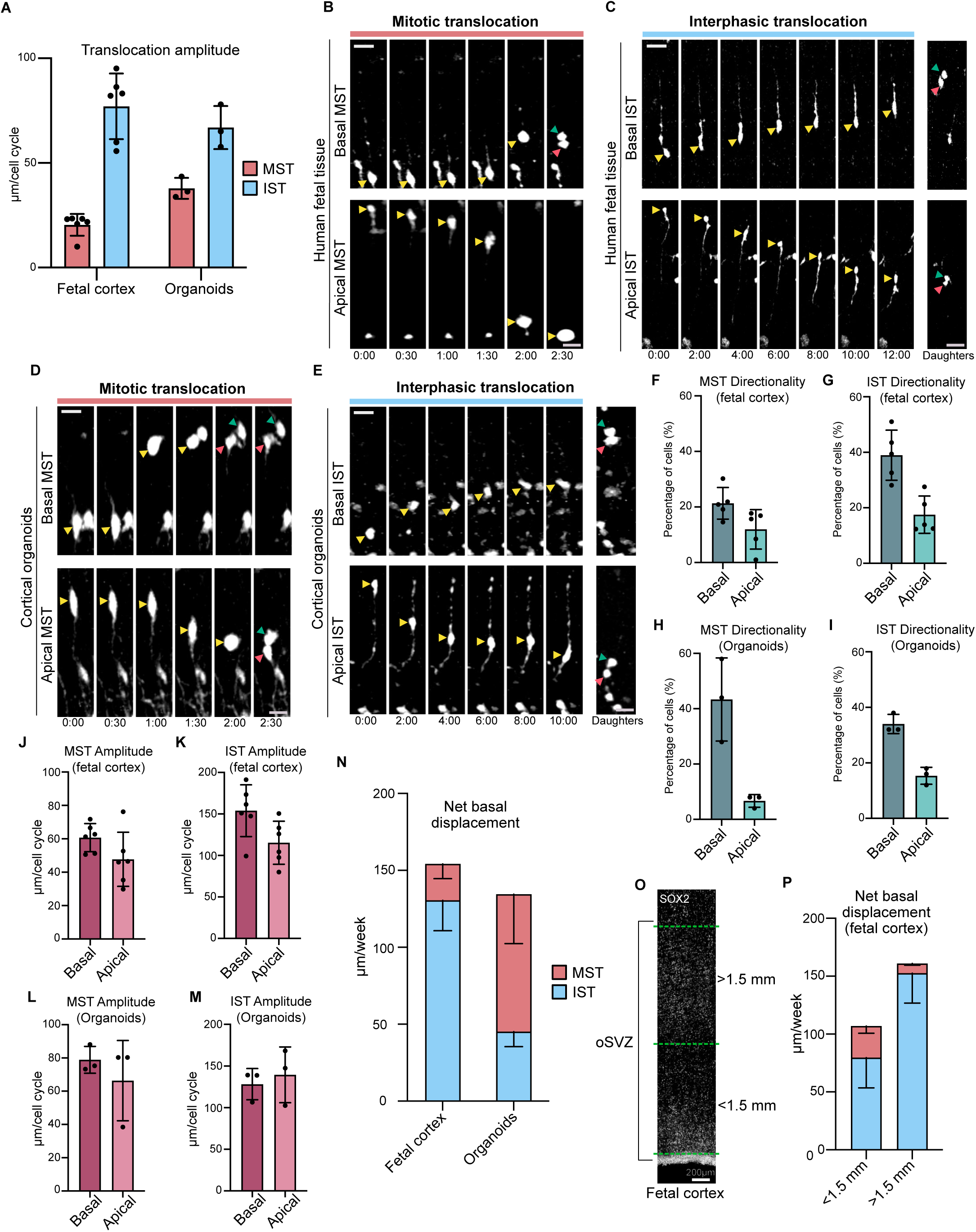
IST contributes more than MST to bRG cells colonization of the human fetal neocortex. (A) Quantification of the amplitude of IST and MST in human fetal tissue (N=5 fetal samples, 476 bRG cells, pcw 12-19. N=3 organoid batches, 150 bRG cells, week 7-9) (B) Live imaging of human bRG cells expressing GFP, performing apically or basally directed MST in human fetal tissue (pcw 14). (C) Live imaging of human bRG cells expressing GFP, performing apically or basally directed IST in human fetal tissue (pcw 14). (D) Live imaging of human bRG cells expressing GFP, performing apically or basally directed MST in human cortical organoids (week 8). (E) Live imaging of human bRG cells expressing GFP, performing apically or basally directed IST in human cortical organoids (week 8). (F) Quantification of the directionality of MST in human fetal tissue (N=5 fetal samples, 476 bRG cells, pcw 12-19). (G) Quantification of the directionality of IST in human fetal tissue (N=5 fetal samples, 476 bRG cells, pcw 12-19). (H) Quantification of the directionality of MST in human cortical organoids (N=3 organoid batches, 150 bRG cells, week 7-9). (I) Quantification of the directionality of IST in human cortical organoids (N=3 organoid batches, 150 bRG cells, week 7-9). (J) Quantification of the amplitude of MST, depending on directionality, in human fetal tissue (N=5 fetal samples, 476 bRG cells, pcw 12-19). (K) Quantification of the amplitude of IST, depending on directionality, in human fetal tissue (N=5 fetal samples, 476 bRG cells, pcw 12-19). (L) Quantification of the amplitude of MST, depending on directionality, in human cortical organoids (N=3 organoid batches, 150 bRG cells, week 7-9). (M) Quantification of the amplitude of IST, depending on directionality, in human cortical organoids (N=3 organoid batches, 150 bRG cells, week 7-9). (N) Contribution of MST and IST to the total net basal displacement of bRG cells, in human fetal tissue and in human cortical organoids (N=5 fetal samples, 476 bRG cells, pcw 12-19. N=3 organoid batches, 150 bRG cells, week 7-9). (O) SOX2 immunostaining of a human fetal cortical slices at pcw 15. The apical region and the basal regions of the oSVZ have been separated by a yellow line, 1.5mm from the ventricular surface. |(P) Contribution of MST and IST to the total net basal displacement of bRG cells, in human fetal tissue in the apical and basal oSVZ (above and below 1.5mm from the ventricular surface) (N= 2 fetal samples, 260 bRG cells, pcw 15-19). Yellow arrowheads indicate bRG cell soma, and green and red arrowheads indicate daughter cells. Data are presented as mean values +/− SD. Scale bar = 20 µm. All live imaging montages are in hours:minutes.

A second factor influencing the contribution of IST and MST to bRG cell colonization of the human cortex is their directionality. As previously reported for MST (Betizeau et al., 2013; Coquand et al., 2024), we found both IST and MST to be bidirectional, occurring in the apical and basal directions, in human fetal tissue and cerebral organoids (Figures 5B-E). Translocation was apical in cells with an apical process, and basal in cells with a basal process. When cells were bipolar, IST and MST generally occurred in the thicker process. Quantification of directionality in human fetal tissues revealed that IST and MST were both biased towards the basal side (Figure 5F & G). IST was slightly more polarized than MST (2.2 versus 1.7 basal bias, respectively). In organoids, MST showed a much stronger basal bias than IST (6.6 versus 2.2 basal bias, respectively) (Figure 5H & I). Whether cells translocated apically or basally, they did so with similar amplitudes, in fetal tissues and in organoids (Figure 5J-M). Together, this indicates that IST and MST display a comparable polarity bias towards the basal side in the fetal cortex, but that MST is more basally polarized than IST in organoids.

We next integrated all these results to quantify how much IST and MST contributed to basal dissemination of bRG cells, taking into account their respective frequency, directionality, and amplitude. In human fetal tissues, bRG cells progressed basally by 155 *μ*M per week of gestation (Figure 5N). IST was the strongest contributor to this movement, accounting for 130 *μ*M of movement, while MST only accounted for 25 *μ*M. Therefore, IST contributes to 85% of the total basal translocation of bRG cells in the human developing neocortex. This difference is largely due to the greater amplitude of movement of IST, which occurs throughout the 46-hour cell cycle, rather than to its biased polarity. In cortical organoids, the total net basal displacement of bRG cells was similar to that of fetal tissues (135 *μ*M per week) (Figure 5N). However, unlike in fetal tissue, MST was the strongest contributor to bRG cell basal displacement, representing 66% of the total.

Finally, we asked whether the translocation characteristics of bRG cells were different depending on their position along the apico-basal axis of the human fetal cortex. We separated the oSVZ into two equal parts, defining a lower and an upper oSVZ (below and above 1.5 mm from the ventricle) (Figure 5O). Cells in the lower oSVZ relied less on IST and more on MST than cells in the upper oSVZ (Figure 5P). In this regard, bRG cells in organoids - where the oSVZ is very small - resemble more bRG cells in the lower part of the human fetal oSVZ. Overall, these results show that IST contributes 5.5 times more than MST to basal dissemination of bRG cells in the developing human fetal cortex.

### IST and MST are conserved in glioblastoma cells

The presence of embryonic bRG-like cells has been reported within adult glioblastoma (GBM) samples (Bhaduri et al., 2020; Wang et al., 2020). These cells were proposed to act as a cancer stem cell-like pool, through reactivation or maintenance of their developmental program. Notably, these cells maintain MST, further validating the parallels between bRG cells and this population of GBM cells (Bhaduri et al., 2020). We therefore asked whether GBM cells also underwent IST, and whether the molecular mechanisms that we uncover here for IST and MST also act in these cells. Because GBMs are highly heterogenous in cell composition (Eisenbarth and Wang, 2023), we live imaged a panel of 9 different GBM lines maintained under neural stem cell culture conditions, and scored for the ones where IST or MST were seen in more than 10% of the cells (Xie et al., 2015). The analysis confirmed heterogeneity and revealed that 5/9 lines performed IST, while 4/9 lines cells performed MST (Figures 6A-D). Notably, line U3123 displayed frequencies of IST and MST extremely similar to that of bRG cells in vitro. These cells were furthermore positive for the bRG cell markers SOX2 and HOPX (Figure 6E). These results indicate that IST and MST are indeed properties of certain GBM cells.

**Figure 6.**
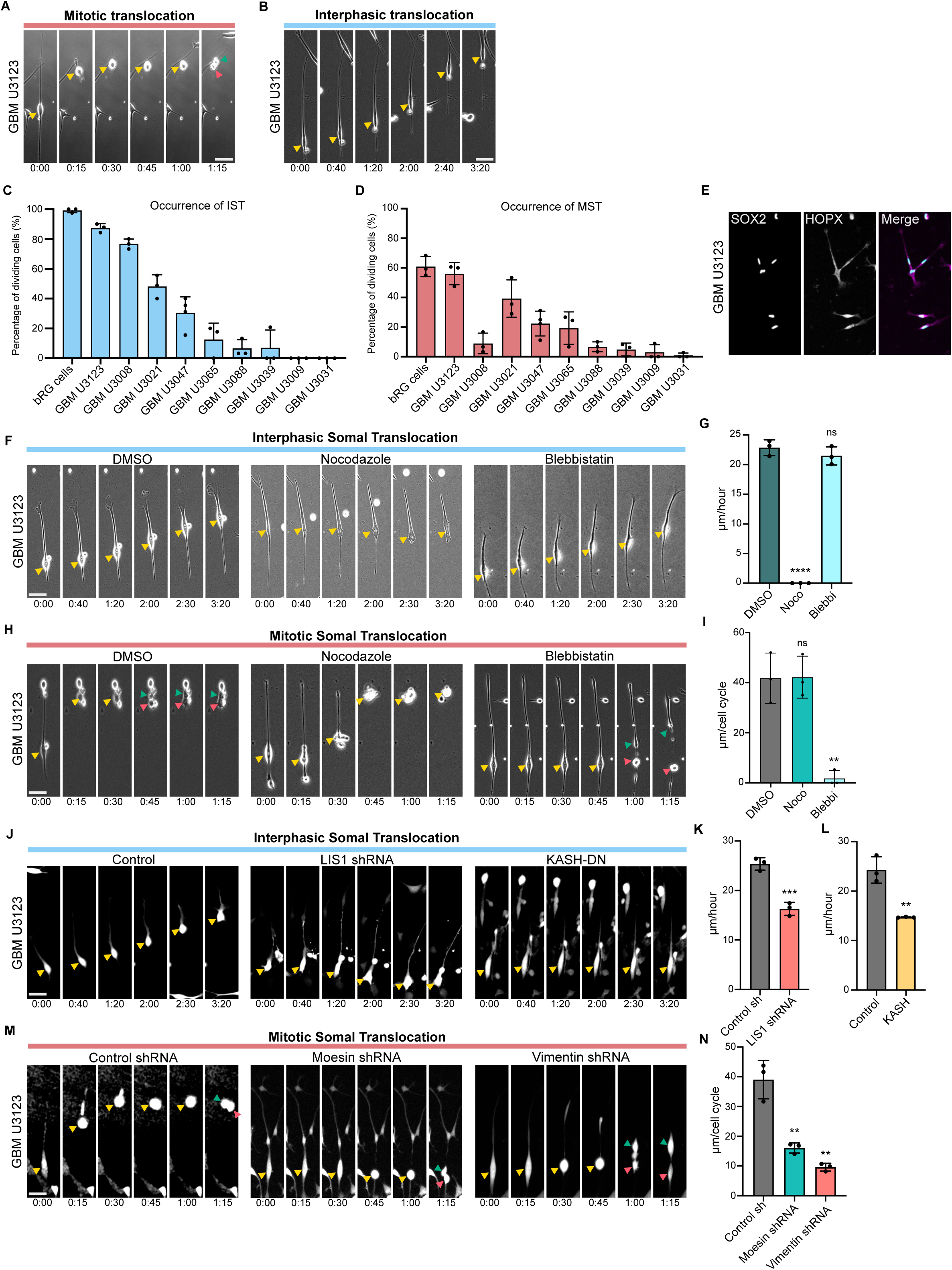
IST and MST are conserved in glioblastoma cells. (A) Live imaging of an *in vitro* GBM cell (line U3123) performing IST. (B) Live imaging of an *in vitro* GBM cell (line U3123) performing MST. (C) Quantification of the fraction of cells performing IST, in 9 GBM lines and compared to *in vitro* bRG cells (N=3 replicates per line, 1130 cells). (D) Quantification of the fraction of cells performing MST, in 9 GBM lines and compared to *in vitro* bRG cells (N=3 replicates per line, 1130 cells). (E) Immunostaining for bRG markers SOX2 and HOPX in U3123 GBM line. (F) Live imaging of U3123 GBM line during interphase, treated with DMSO, nocodazole (1*μ*M) or blebbistatin (10*μ*M). (G) Quantification of IST amplitude following treatment with DMSO, nocodazole (1*μ*M) or blebbistatin (10*μ*M) in U3123 GBM line (N=3 experiments, 274 GBM cells). (H) Live imaging of U3123 GBM line during mitosis, treated with DMSO, nocodazole (1*μ*M) or blebbistatin (10*μ*M). (I) Quantification of MST amplitude following treatment with DMSO, nocodazole (1*μ*M) or blebbistatin (10*μ*M) in U3123 GBM line (N=3 experiments, 274 GBM cells). (J) Live imaging of interphasic U3123 GBM cells expressing control, LIS1 shRNA or KASH dominant negative constructs. (K) Quantification of IST amplitude in U3123 GBM cells expressing control or LIS1 shRNA constructs. (N=3 experiments, 359 GBM cells). (L) Quantification of IST amplitude in U3123 GBM cells expressing control or KASH dominant negative constructs (N=3 experiments, 300 GBM cells). (M) Live imaging of mitotic U3123 GBM cells expressing control, Moesin or Vimentin shRNA constructs. (N) Quantification of MST amplitude U3123 GBM cells expressing control, Moesin or Vimentin shRNA (N=3 experiments, 149 GBM cells). Yellow arrowheads indicate bRG cell soma, and green and red arrowheads indicate daughter cells. Data are presented as mean values +/− SD. Scale bar = 20 µm. All live imaging montages are in hours:minutes. **p<0,01; ***p<0,001; ****p<0,0001, ns: non-significant by two-tailed unpaired t-tests.

We next used line U3123 to test if the molecular mechanisms of IST and MST were conserved in GBM cells. Incubation of U3123 with Nocodazol completely abolished IST, while Blebbistatin had no effect (Figure 6F & G). Conversely, incubation with Blebbistatin completely abolished MST, while Nocodazol had no effect (Figure 6H & I). Therefore, as for bRG cells, IST is microtubule-dependent, and MST is actomyosin-dependent in GBM cells. We then asked if the underlying molecular mechanisms for both of these movements were also conserved. shRNA-mediated knockdown of LIS1 indeed altered IST without affecting MST in GBM cells (Figures 6J & K). Likewise, expression of the LINC complex dominant negative KASH construct reduced IST in the cells (Figures 6J & L). Conversely, knockdown of Moesin and Vimentin in GBM cells impaired MST, without affecting IST (Figures 6M & N). Together, these results indicate that the molecular mechanisms of IST and MST are conserved between fetal bRG cells and GBM cells.

## Discussion

Our work provides a descriptive and mechanistic study of basal radial glial cell colonization of the human fetal neocortex. We identify two distinct modes of translocation: IST, occurring in interphase in a microtubule-dependent manner, and MST, occurring during mitosis, relying on the actin cytoskeleton. We demonstrate that IST is controlled by dynein and its activator LIS1 which are recruited to the nuclear envelope by the LINC complex, while MST is driven by the mitotic cell rounding pathway, through ERM proteins, Vimentin and actomyosin contractility. IST contributes five times more than MST to the dissemination of bRG cells in the human fetal cortex, for a total movement of 0,67 mm per month of gestation. We furthermore show that IST is altered in lissencephalic patients, and that bRG-like glioblastoma cells also utilize IST and MST for their movement. This work therefore identifies a biphasic movement for neural stem cell colonization of the human fetal cortex, with important implications in pathological contexts.

Similarly to apical nuclear migration in aRG cells and nucleokinesis in migrating neurons, IST in bRG cells depends on the dynein motor. We show here that the LINC complex-dependent pathway for dynein recruitment to the nuclear envelope is the same as in neurons but different from aRG cells, where dynein is recruited to the nuclear pore complex. Therefore, IST is molecularly closer to neuronal migration than it is to INM, even though bRG cells are transcriptionally much closer to aRG cells than to neurons (Qian et al., 2024; Wang et al., 2024). This is likely because apical nuclear migration in aRG cells is a G2-specific mechanism, while IST and neuronal migration are not. A notable difference between neurons and bRG cells is that neuronal migration is dependent on actomyosin while IST is not. This may reflect a prevalent role for actomyosin in neuronal leading-edge progression, while bRG cells have pre-extended static processes that do not need to grow during translocation.

We show that MST is a mitotic spindle translocation event that occurs after nuclear envelope breakdown and is driven by the process of mitotic cell rounding. This well-described pathway induces an important stiffening of the cell cortex, enabling adherent cells to round up for proper chromosomal segregation. RhoA-ROCK-Myosin II, the ERM proteins - which crosslink the actomyosin network to the plasma membrane - and vimentin - which underlies the actin network and increases stiffness - all participate in MST. Mitotic cell rounding is therefore the force that drives MST, and we propose that two phenomena transform this force into somal movement. First, focal adhesions are destabilized in the soma, a phenomenon well-known to occur during mitotic cell rounding, which releases the soma from the ECM. Second, the large intracellular tension that builds up in the soma is dissipated into the basal process of bRG cells, which leads to its engulfment into the basal process. Why some cells do not perform MST and remain static is unclear, but this may be due to incomplete focal adhesion disassembly or reduced intracellular tension.

While bRG cell translocation is the mechanism by which they colonize the neocortex, the actual role of this process remains unclear. bRG cells were shown to undergo more direct or indirect divisions depending on their apicobasal position within the tissue, so a role for IST and MST may be to position bRG cells into different microenvironments that would influence their output (Coquand et al., 2024). This question is however highly challenging to address, as differences in cell fate decisions are observed between regions distant by hundreds of *μ*M, which bRG cells take weeks to cover in the fetal cortex.

bRG cell translocation might also contribute to neuronal positioning. We indeed show here that, as neurogenesis progresses, bRG cells advance more and more basally into the neocortex. Consequently, they generate newborn neurons further and further away from the ventricular surface. This reduces the distance neurons must travel to reach the top of the cortical plate, as compared to neurons born from aRG cells. In this regard, it is interesting to note that the pathway controlling IST and neuronal migration is largely the same, and that loss of function of the underlying gene, for example dynein, will affect both bRG translocation and neuronal migration. The neuronal positioning phenotype associated with these factors must therefore be seen as the addition of bRG cell and neuronal defects. This is particularly relevant in the case of lissencephaly, a neuronal positioning disease. We show here that in LIS1-mutated patient organoids, IST is severely affected. In these patients, neurons will therefore be born farther away from their final location, amplifying the neuronal migration defects.

A last possible function for bRG cell somal translocation could be to avoid the jamming of these cells. bRG cells indeed have a very high self-amplification capacity, leading to their rapid expansion between weeks 12 and 20 of human fetal gestation (Coquand et al., 2024). Massive amounts of bRG cells might therefore accumulate in the same location, close to the ventricles where they are born. We speculate that alteration of their ability to translocate could lead to heterotopias, which are abnormal clusters of cells that end up in the white matter postnatally. Genes associated with heterotopia are strongly associated with the cytoskeleton, with actin-related genes - such as FLNA - that may affect MST, and microtubule-related genes - such as MAP1B - that may affect IST (Romero et al., 2018).

We show here that bRG cells translocate bidirectionally, for a net basal displacement of 0,67 mm per month of human fetal gestation. On top of this active motion, a passive displacement of bRG cells will also occur as a consequence of cell proliferation and tissue growth. Indeed, as bRG cells actively translocate, they expand the oSVZ size and displace its basal boundary. The massive proliferation between these basal cells and the ventricular surface will further push them away, increasing their displacement. Therefore, the combination of oSVZ proliferation and somal translocation together account for the total basal displacement of bRG cells. Our works provides a mechanistic understanding of oSVZ growth during human fetal development, a core feature of neocortical expansion during evolution, and a process likely to be altered in several cortical malformations.

---

**Supplementary Figure 1.**
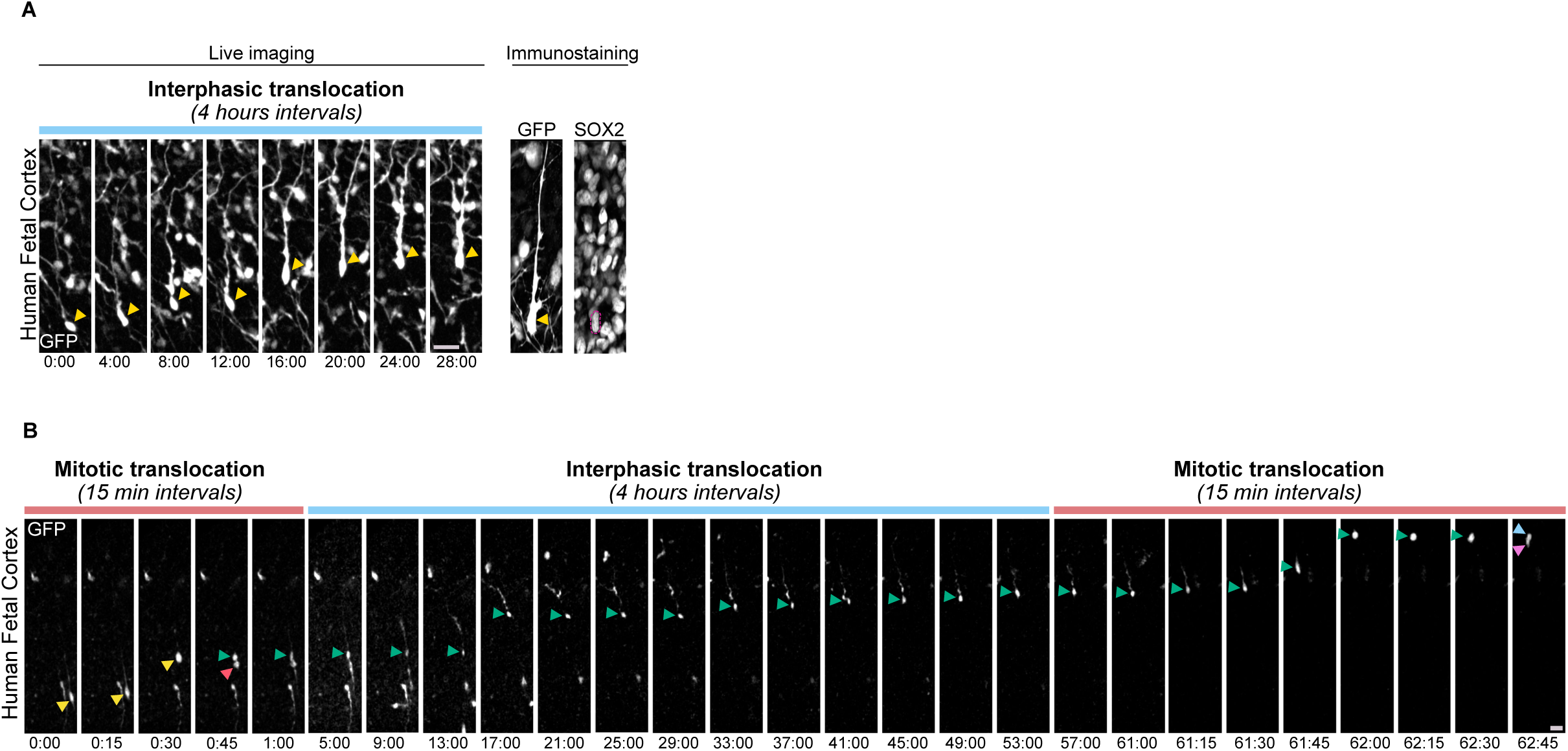
Fate of IST-performing cells and long-term imaging of translocation. (A) Live-fix correlative imaging revealing a bRG cell undergoing interphasic somal translocation and positive for SOX2 in human fetal tissue at pcw 15. Yellow arrowheads indicate bRG cell soma, and green and red arrowheads indicate daughter cells. (B) 4-day live imaging of a human bRG cell expressing GFP in fetal cortex at pcw 16. Following MST and division, a self-renewing bRG daughter cell undergoes IST, followed by as second MST and division. Yellow arrowhead indicates bRG cell soma, and green and red arrowheads indicate daughter cells. Green arrowhead bRG daughter is followed during its entire cell cycle. Blue and pink arrowheads indicate daughter cells. Scale bar = 20 µm. All live imaging montages are in hours:minutes.

**Supplementary Figure 2.**
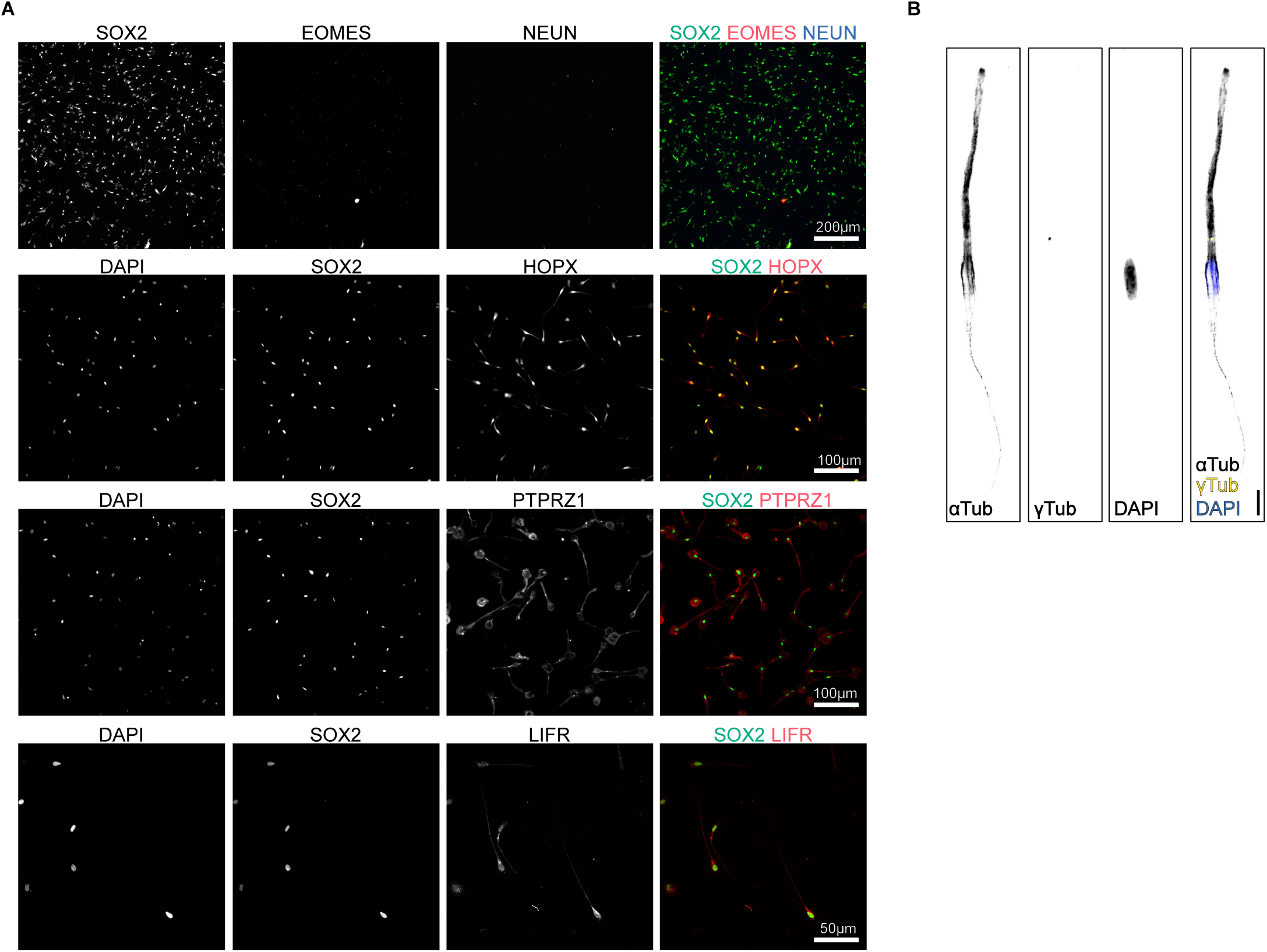
Fate of *in vitro* bRG cells and microtubule organization. (A) Cell fate analysis of *in vitro* bRG cells. Cells were stained for IP marker EOMES, Neuronal marker (NEUN) and bRG markers SOX2, HOPX, PTPRZ1 and LIFR. (B) *In vitro* bRG cells stained for ⍺-Tubulin to visualize the microtubule cage and %-Tubulin to visualize the centrosome. Scale bar = 20 µm

**Supplementary Figure 3.**
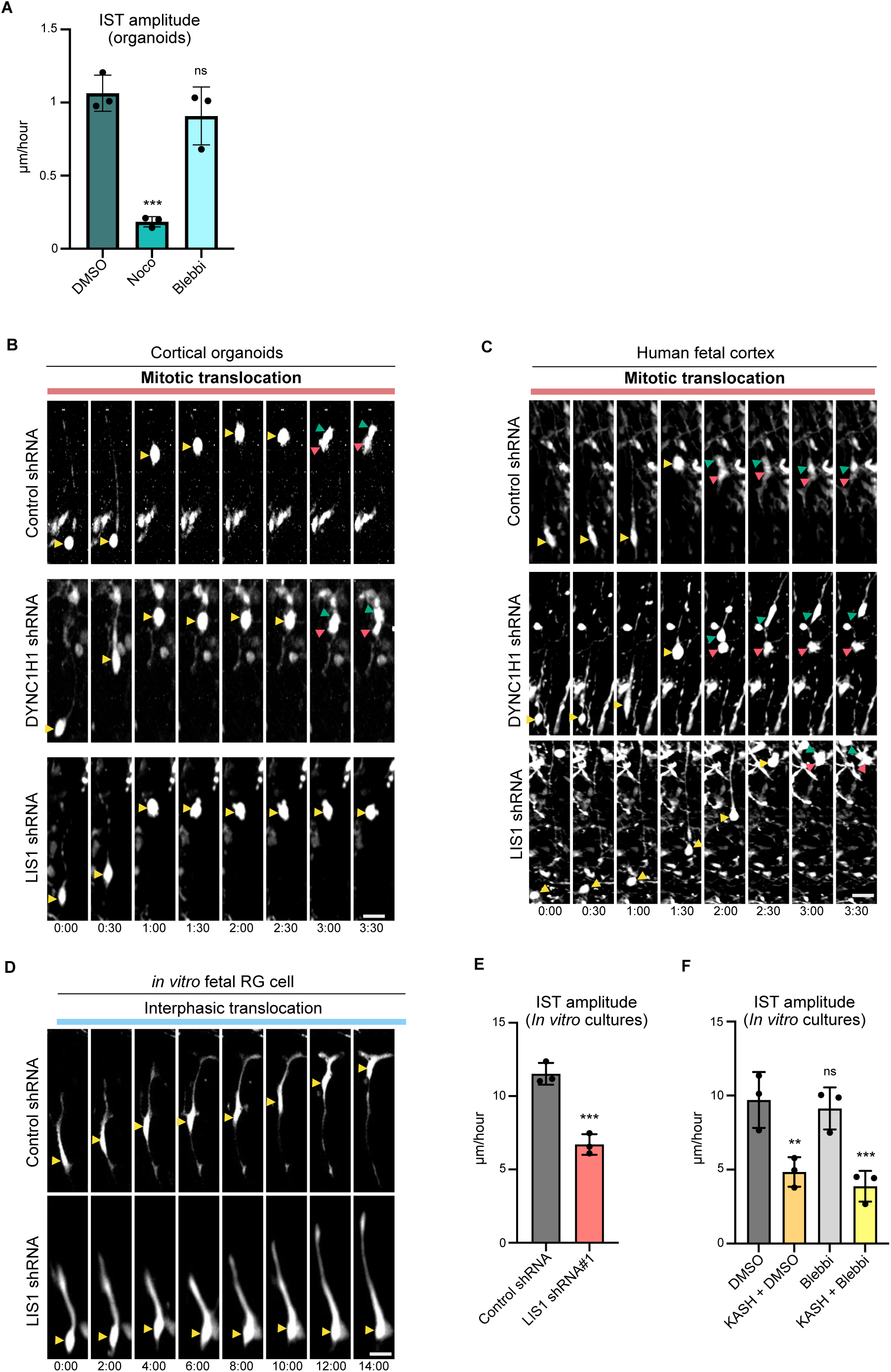
Drug treatments and live imaging examples. (A) Quantification of IST amplitude in bRG cells following treatment with DMSO, nocodazole (1*μ*M) or blebbistatin (10*μ*M) in cortical organoids (N=3 organoid batches, 329 bRG cells, weeks 9-12). (B) Live imaging of mitotic human bRG cells expressing control, DYNC1H1 or LIS1 shRNA constructs in human cortical organoids (week 8-11). shRNA plasmids co-express GFP. (C) Live imaging of mitotic human bRG cells expressing control, DYNC1H1 or LIS1 shRNA constructs in human fetal tissue (pcw 16-20). (D) Live imaging of *in vitro* interphasic human bRG cells expressing control or LIS1 shRNA constructs. (E) Quantification of IST amplitude in *in vitro* interphasic human bRG cells expressing control or LIS1 shRNA constructs (N=3 experiments, 520 bRG cells). (F) Quantification of IST amplitude in *in vitro* interphasic human bRG cells expressing control or KASH constructs, in the presence of DMSO or blebbistatin (10*μ*M) (N=3 experiments, 1198 bRG cells). Yellow arrowheads indicate bRG cell soma, and green and red arrowheads indicate daughter cells. Data are presented as mean values +/− SD. Scale bar = 20 µm. All live imaging montages are in hours:minutes. **p<0,01; ***p<0,001, ns: non-significant by two-tailed unpaired t-tests.

## Materials and Methods

### Statistics and reproducibility

Quantitative data are described as mean ± s.d. for N ≥ 3, except for some human fetal tissue data which are described as N = 2, without s.d. No data were excluded from the analyses and the experiments were not randomized. No statistical method was used to predetermine sample size. The investigators were not blinded to allocation during experiments and outcome assessment. Statistical analysis was performed using a two-tailed unpaired Student’s t-test using GraphPad Prism 10 software (GraphPad Software). Data distribution was assumed to be normal. P values lower than 0.05 were considered statistically significant. The figures were created using Affinity Designer 1.10.

### Virus production

We employed the HEK-Phoenix-GP cell line obtained from ATCC (CRL-3215), which stably expresses the packaging enzymes GAL and POL. Cells were seeded in 4×T300 flasks at a 1:20 dilution and cultured for three days in DMEM-GlutaMax medium supplemented with 10% fetal bovine serum (FBS) (50 ml per flask) until they reached approximately 70% confluence. On the third day, the cells were transfected with an envelope VSVG plasmid and a transfer plasmid (either CAG-GFP or MSCV-IRES-GFP) using Lipofectamine 2000. For this, the plasmids were mixed in 14.4 ml of OptiMEM medium (24 µg of envelope plasmid and 66 µg of transfer plasmid). Concurrently, 450 µl of Lipofectamine 2000 was diluted in 14.4 ml of OptiMEM medium and allowed to incubate for 5 minutes at room temperature. The DNA mixture was then combined with the Lipofectamine solution and incubated for an additional 30 minutes at room temperature. Meanwhile, the medium in each T300 flask was replaced with 30 ml of DMEM-GlutaMax (without FBS). Following the incubation, 3.6 ml of the DNA-Lipofectamine mixture was added to each flask, and the cells were incubated for 5 hours at 37°C. For lentiviral production, we used HEK T293 cells. Cells were plated in 3xT150 (dilution 1:20) and grown for 3 days to reach 70% confluency in DMEM-GlutaMax medium, 10% foetal bovine serum (FBS) (30 ml per flask). On day 3, cells were transfected with PMD2G (an envelope expressing VSVG), PMAX2 (a second packaging vector) and the transfer plasmid. Subsequently, the retrovirus and lentivirus flasks were moved to an L3 laboratory where the medium was replaced with 30 ml of fresh DMEM-GlutaMax containing 10% FBS. On the fifth day, the medium was harvested into 50 ml tubes, replaced with 30 ml of fresh medium, and stored at 4°C. On the sixth day, the medium was again collected, pooled with the day 5 samples, and centrifuged at 1,300 rpm for 5 minutes at 4°C to pellet cell debris. The supernatant was then filtered through a 0.22 µm filter unit and distributed into six Ultra-Clear tubes (Beckman Coulter, 344058). These tubes were ultracentrifuged at 31,000g for 90 minutes at 4°C. The supernatant was discarded, and the retrovirus-containing pellet was washed several times with PBS and transferred to a new Ultra-Clear tube. A final ultracentrifugation step was performed (31,000g for 90 minutes at 4°C), after which the supernatant was carefully removed, and the retrovirus pellet was resuspended in 1 ml of DMEM-F12 medium. The suspension was aliquoted (50 to 100 µl per aliquot) and stored at −80°C. The viral titer of the preparation was assessed by infecting standard HEK cells at various dilutions, and the proportion of GFP-positive cells was determined using fluorescence-activated cell sorting (FACS).

### Human iPS cell culture

The feeder-independent induced pluripotent stem (iPS) cell lines used for this study were obtained through collaborations with the laboratory of Silvia Cappello (Max Planck Institute of Psychiatry), Julia Ladewig (Hector Institute for Translational Brain Research) and Laurent David (University of Nantes). HMGU1 cells were reprogrammed from NuFF3-RQ human newborn foreskin feeder fibroblasts (GSC-3404, Global Stel) (Kyrousi et al., 2021). The f71.002 wild-type line - from a female donor (Gaignerie et al., 2018) - was a kind gift from Laurent David (iPSC core facility from Nantes Université, supported by IBiSA and Biogenouest). iPS cells were cultivated as colonies on vitronectin-coated B3 dishes, using StemFlex medium (Thermo Fisher Scientific). Colonies were cleaned every two days under a binocular stereo microscope (Lynx EVO, Vision Engineering), by manually removing differentiated cells with a needle. Two independent LIS1-mutated patient derived iPS cells were obtained from the laboratory of Julia Ladewig. Patient #1, LIS1-severe 1, 4-year-old female donor, c.1002+1G>T and patient #2 LIS1-severe 2, 18-year-old female donor, c.531G>C LIS1-mutation. Each patient specific mutation was confirmed through Sanger sequencing and quality control by the laboratory of origin (Zillich et al., 2022).

### Generation of Cortical Organoids

Cerebral organoids were generated from human induced pluripotent stem (iPS) cells following an adapted protocol from previously published (Qian et al., 2016; Sloan et al., 2018). Day 0 to Day 4: iPS cell colonies were detached using pre-warmed collagenase (1 mg/ml) for 45 minutes at 37°C. The floating colonies were transferred using a cut pipette tip into a 15 ml tube and washed with Medium 1 (DMEM-F12 without phenol red, 20% KnockOut Serum Replacement (KSR), 1× GlutaMAX, 1× MEM Non-Essential Amino Acids (MEM-NEAA), 1× 2-Mercaptoethanol, Penicillin/Streptomycin, 2 µM Dorsomorphin, 2 µM A-83). The colonies were then transferred into ultra-low attachment plates with 3 ml of Medium 1 and cultured at 37°C in a 5% CO2 atmosphere. Day 5 to Day 6: Half of the Medium 1 was replaced daily with Medium 2 (DMEM-F12 without phenol red, 1× N2 supplement, 1× GlutaMAX, 1× MEM-NEAA, Penicillin/Streptomycin, 1 µM CHIR-99021, 1 µM SB-431542). Day 7 to Day 14: On Day 7, embryoid bodies (EBs) were embedded in Matrigel diluted in Medium 2 at a ratio of 2:1. The Matrigel-EB mixture was spread into regular 6-well plates and incubated at 37°C for 30 minutes to solidify (10-20 EBs per well). Medium 2 was gently added to the well without disturbing the Matrigel patch. On Day 14, the Matrigel-EB mixture was mechanically dissociated using a 5 ml pipette and transferred into a 15 ml tube for gentle washing. Organoids were then suspended in Medium 3 (DMEM-F12 without phenol red, 1× N2 supplement, 1× B27 supplement (with Vitamin A), 1×GlutaMAX, 1×MEM-NEAA, 1× 2-Mercaptoethanol, Penicillin/Streptomycin, 2.5 µg/ml insulin) and cultured in ultra-low attachment 6-well plates under agitation at 100 rpm using a Digital Orbital Shaker (DOS-10M from ELMI). Day 35 to Day 84: Starting from Day 35, Medium 3 was supplemented every two days with diluted Matrigel (1:100) to support the growth and development of the cerebral organoids.

### Infection of human fetal cortex and cerebral organoids

Fresh human fetal prefrontal cortex samples were obtained from autopsies conducted at Robert Debré Hospital and Necker-Enfants Malades Hospital in Paris. These tissues were derived from spontaneous miscarriages or pregnancy terminations due to organ malformations. A sample of the pre-frontal cortex was excised from one hemisphere and transported on ice to the laboratory. The tissue was subdivided into smaller pieces and embedded in 4% low-gelling agarose (Sigma), which was dissolved in DMEM-F12. Cerebral organoids (8–12 weeks) were embedded in 3% low-gelling agarose. Both types of tissue were sectioned into 300-µm-thick slices using a Leica VT1200S vibratome in ice-cold DMEM-F12. The slices were infected with a GFP-coding retrovirus diluted in DMEM-F12. After a 2-hour incubation period, the slices were washed three times with DMEM-F12 and cultured on Millicell cell culture inserts (Merck) in cortical culture medium (DMEM-F12 supplemented with B27, N2, 10 ng/ml fibroblast growth factor (FGF), 10 ng/ml epidermal growth factor (EGF), 5% foetal bovine serum, and 5% normal horse serum).

### Live imaging of cortical slices

Fetal tissue (96 hours post-infection) and organoid slices (48 hours post-infection) were transferred to a 35 mm FluoroDish (WPI) containing 1 ml of cortical culture medium. 48 to 96-hour live imaging was performed using a spinning disk-wide microscope equipped with a Yokogawa CSU-W1 scanner unit to enhance the field of view and resolution. The microscope was equipped with a high working distance (WD 6.9-8.2 mm) ×20 Plan Fluor ELWD NA 0.45 dry objective (Nikon) and a Prime95B SCMOS camera. Z-stacks ranging from 100 to 110 µm were captured at 5 µm intervals, and maximum projections were generated. Videos were assembled and processed using Fiji ImageJ, with maximum projections, background subtraction, median filtering, stack registration, and rotation adjustments applied.

### Isolation and culture of human fetal radial glial cells

Fresh human fetal prefrontal cortex tissue was washed with culture medium (DMEM-F12, D-Glucose 2.9 mg/mL, Penicillin/Streptomycin 5U/mL, Amphotericin 250ng/mL) and replaced by culture medium supplemented with B27 (-vitamin A), FGF 20 ng/mL and EGF 20 ng/mL. The tissue was then disassociated by mechanical trituration with a pipette. The suspension was spun down at 1200rpm for 3mins and the pellet suspended into 1mL of supplemented culture media. After viability was checked (70-90%), the cells were plated into 6 well plates coated with 0.1 mg/mL Poly-D-Lysine and 1.5% Fibronectin, in 2mL of culture media at a density of 2.000.000 cells per well. The cells were incubated at 37°C, 5% C02. The culture medium was replaced the day after and every two days afterwards to remove floating cells and debris. 2-3 weeks later, depending on the tissue stage, radial glia cells were detached with pre-warmed accutase for 10mins at 37°C, 5% C02 and replated into novel pre-coated plates. Cells were checked for fate markers at each consecutive passage by fixing with 4% PFA and immunostaining for fate markers.

### Infection of *in-vitro* radial glia cells

Radial Glia cells were plated onto 6-well glass bottom plates (IBL P06-1.5H-N), coated with the above-mentioned substrate. Radial Glia cells were transduced with lentiviral and retroviral particles for 2 hours at 37°C, 5% C02. After washing with non-supplemented media, 2mL of supplemented media was added per well. After 24 to 48 hours of expression of the different constructs, the cells were imaged on a TI2-E inverted video-microscope equipped with a photometric Kinetix sCMOS camera, and each position was imaged every 5 minutes in GFP or RFP with a 10X Plan Fluor, NA: 0.20, WD 15.20 dry objective, using a perfect focus (PFS) mode. To reduce phototoxicity, the laser power was used at the lowest power possible, and the images were binned to increase the fluorescent signal.

### Drug treatments

To inhibit actomyosin contractility and to depolymerise microtubules, *in-vitro* radial glia cells were plated in 6 well plates and brought to a TI2-E inverted video-microscope with a photometric Kinetix sCMOS camera. Each position was imaged every 5 minutes in phase contrast with a 10X Plan Fluor, NA: 0.20, WD 15.20 dry objective, using a perfect focus (PFS) mode. At the start of the movie culture media was supplemented with10µM of blebbistatin or 1µM of Nocodazole, or with a corresponding volume of DMSO.

### Culture of Glioblastoma Cell Lines

Patient-derived glioblastoma cells were acquired from the Human Glioblastoma Cell Culture resource (www.hgcc.se) at the Dept. of Immunology, Genetics and Pathology, Uppsala University, Uppsala, Sweden and are a part of collaboration through a Material Transfer Agreement between Institute Curie and Uppsala University (Xie et al., 2015). U3008, U3009, U3017, U3021, U3031, U3039, U3047, U3065, U3088, U3123 cell lines from the HGCC biobank were cultured in cell culture plastic dishes coated with Matrigel in culture medium containing 50% DMEM-F12 and 50% Neurobasal Medium, Penicillin/Streptomycin 1U/mL, B27 (-vitamin A), EGF 10ng/ml and FGF 10ng/ml. At confluency for imaging or for maintenance, cells were detached using pre-warmed accutase and replated in Matrigel coated dishes. Treatment of these cells with nocodazole (1µM), blebbistatin (10µM) or infection with different viral particles and subsequent imaging was done as described for in vitro radial glial cells.

### Expression constructs and antibodies

The following plasmids were used in this study: MSCV-IRES-GFP (Tannishtha Reya, Addgene 20672); VSVG (a gift from P. Benaroch), Human EZR shRNA (TF308420, Origene), Human STK10 shRNA (TF320540, Origene), Human SLK shRNA (TG320620, Origene), Human DYNC1H1 shRNA (TL313335, Origene), Human RDX shRNA (TL309884, Origene), Human MSN shRNA (TL311375, Origene), Human ECT2 shRNA (TL304854, Origene), Human VIM shRNA (TL308419, Origene), Human PAFAH1B1 (LIS1) shRNA (TL310628, Origene), Dominant Negative KASH. Antibodies used in this study were mouse anti-SOX2 (Abcam Ab79351, 1/500), sheep anti-EOMES (R&D Sytems AF6166, 1/500), rabbit anti-NEUN (Abcam Ab177487, 1/500), chicken anti-GFP (Abcam Ab13970, 1/500), mouse anti-pVimentin (Abcam Ab22651, 1/1000), rabbit anti-NeuroD2 (Abcam, ab104430, 1/500), rabbit anti-HOPX (Proteintech, 11419-1-AP, 1/500), rabbit anti-NESPRIN2 (Abcam ab204308, 1/500), rabbit anti-PH3 (Abcam ab47297, 1/2000), mouse anti-gamma Tubulin (Sigma-Aldrich T5326, 1/1000), rabbit anti-pERM (Cell Signalling Technology, 3141S), mouse anti-LIFR (Abcam 89792, 1/500), rabbit anti-PTPRZ1 (Sigma HPA015103 Atlas antibodies, 1/500). Secondary antibodies used were: Donkey Anti-Sheep IgG H&L (Alexa Fluor 405) Abcam ab175676; DyLight 405 AffiniPure Donkey Anti-Mouse IgG (H+L) Jackson ImmunoResearch 715-475-150; DyLight 405 AffiniPure Donkey Anti-Rabbit IgG (H+L) Jackson ImmunoResearch 711-475-152; Alexa Fluor 488 AffiniPure Donkey Anti-Rabbit IgG (H+L) Jackson ImmunoResearch 711-545-152; Donkey anti-Rabbit IgG (H+L) Highly Cross-Adsorbed Secondary Antibody, Alexa Fluor Plus 488 Thermo Fisher A32790; Donkey anti-Chicken IgY (H+L) Highly Cross-Adsorbed Secondary Antibody, Alexa Fluor 488 Thermo Fisher A78948; Alexa Fluor 488 AffiniPure Donkey Anti-Chicken IgY (IgG) (H+L) Jackson ImmunoResearch 703-545-155; Alexa Fluor 488 AffiniPure Donkey Anti-Mouse IgG (H+L) Jackson ImmunoResearch 715-545-150; Donkey anti-Mouse IgG (H+L) Highly Cross-Adsorbed Secondary Antibody, Alexa Fluor Plus 488 Thermo Fisher A32766; Donkey anti-Sheep IgG (H+L) Cross-Adsorbed Secondary Antibody, Alexa Fluor 568 Thermo Fisher A21099; Cy3 AffiniPure Donkey Anti-Sheep IgG (H+L) Jackson ImmunoResearch 713-165-147; Donkey anti-Mouse IgG (H+L) Highly Cross-Adsorbed Secondary Antibody, Alexa Fluor 568 Thermo Fisher A10037; Cy3 AffiniPure Donkey Anti-Mouse IgG (H+L) Jackson ImmunoResearch 715-165-150; Donkey anti-Rabbit IgG (H+L) Highly Cross-Adsorbed Secondary Antibody, Alexa Fluor 568 Thermo Fisher A10042; Cy3 AffiniPure Donkey Anti-Rabbit IgG (H+L) Jackson ImmunoResearch 711-165-152; Donkey anti-Rabbit IgG (H+L) Highly Cross-Adsorbed Secondary Antibody, Alexa Fluor Plus 647 Thermo Fisher A32795; Alexa Fluor 647 AffiniPure Donkey Anti-Rabbit IgG (H+L) Jackson ImmunoResearch 715-605-152; Donkey anti-Goat IgG (H+L) Highly Cross-Adsorbed Secondary Antibody, Alexa Fluor Plus 647 Thermo Fisher A32849; Alexa Fluor 647 AffiniPure Donkey Anti-Goat IgG (H+L) Jackson ImmunoResearch 705-605-003; Alexa Fluor 647 AffiniPure Donkey Anti-Mouse IgG (H+L) Jackson ImmunoResearch 715-605-150.

## References

1. Allen, D.E., Donohue, K.C., Cadwell, C.R., Shin, D., Keefe, M.G., Sohal, V.S., Nowakowski, T.J., 2022. Fate mapping of neural stem cell niches reveals distinct origins of human cortical astrocytes. Science 376, 1441–1446. 10.1126/science.abm5224

2. Andrews, M.G., Siebert, C., Wang, L., White, M.L., Ross, J., Morales, R., Donnay, M., Bamfonga, G., Mukhtar, T., McKinney, A.A., Gemenes, K., Wang, S., Bi, Q., Crouch, E.E., Parikshak, N., Panagiotakos, G., Huang, E., Bhaduri, A., Kriegstein, A.R., 2023. LIF signaling regulates outer radial glial to interneuron fate during human cortical development. Cell Stem Cell 30, 1382–1391.e5. 10.1016/j.stem.2023.08.009

3. Andrews, M.G., Subramanian, L., Kriegstein, A.R., 2020. mTOR signaling regulates the morphology and migration of outer radial glia in developing human cortex. eLife 9, e58737. 10.7554/eLife.58737

4. Baffet, A.D., Hu, D.J., Vallee, R.B., 2015. Cdk1 Activates Pre-mitotic Nuclear Envelope Dynein Recruitment and Apical Nuclear Migration in Neural Stem Cells. Dev. Cell 33, 703–716. 10.1016/j.devcel.2015.04.022

5. Betizeau, M., Cortay, V., Patti, D., Pfister, S., Gautier, E., Bellemin-Ménard, A., Afanassieff, M., Huissoud, C., Douglas, R.J., Kennedy, H., Dehay, C., 2013. Precursor Diversity and Complexity of Lineage Relationships in the Outer Subventricular Zone of the Primate. Neuron 80, 442–457. 10.1016/j.neuron.2013.09.032

6. Bhaduri, A., Di Lullo, E., Jung, D., Müller, S., Crouch, E.E., Espinosa, C.S., Ozawa, T., Alvarado, B., Spatazza, J., Cadwell, C.R., Wilkins, G., Velmeshev, D., Liu, S.J., Malatesta, M., Andrews, M.G., Mostajo-Radji, M.A., Huang, E.J., Nowakowski, T.J., Lim, D.A., Diaz, A., Raleigh, D.R., Kriegstein, A.R., 2020. Outer Radial Glia-like Cancer Stem Cells Contribute to Heterogeneity of Glioblastoma. Cell Stem Cell 26, 48–63.e6. 10.1016/j.stem.2019.11.015

7. Coquand, L., Brunet Avalos, C., Macé, A.-S., Farcy, S., Di Cicco, A., Lampic, M., Wimmer, R., Bessières, B., Attie-Bitach, T., Fraisier, V., Sens, P., Guimiot, F., Brault, J.-B., Baffet, A.D., 2024. A cell fate decision map reveals abundant direct neurogenesis bypassing intermediate progenitors in the human developing neocortex. Nat. Cell Biol. 26, 698–709. 10.1038/s41556-024-01393-z

8. Delgado, R.N., Allen, D.E., Keefe, M.G., Mancia Leon, W.R., Ziffra, R.S., Crouch, E.E., Alvarez-Buylla, A., Nowakowski, T.J., 2022. Individual human cortical progenitors can produce excitatory and inhibitory neurons. Nature 601, 397–403. 10.1038/s41586-021-04230-7

9. Del-Valle-Anton, L., Amin, S., Cimino, D., Neuhaus, F., Dvoretskova, E., Fernández, V., Babal, Y.K., Garcia-Frigola, C., Prieto-Colomina, A., Murcia-Ramón, R., Nomura, Y., Cárdenas, A., Feng, C., Moreno-Bravo, J.A., Götz, M., Mayer, C., Borrell, V., 2024. Multiple parallel cell lineages in the developing mammalian cerebral cortex. Sci. Adv. 10, eadn9998. 10.1126/sciadv.adn9998

10. Duarte, S., Viedma-Poyatos, Á., Navarro-Carrasco, E., Martínez, A.E., Pajares, M.A., Pérez-Sala, D., 2019. Vimentin filaments interact with the actin cortex in mitosis allowing normal cell division. Nat. Commun. 10, 4200. 10.1038/s41467-019-12029-4

11. Eisenbarth, D., Wang, Y.A., 2023. Glioblastoma heterogeneity at single cell resolution. Oncogene 42, 2155–2165. 10.1038/s41388-023-02738-y

12. Fernández, V., Borrell, V., 2023. Developmental mechanisms of gyrification. Curr. Opin. Neurobiol. 80, 102711. 10.1016/j.conb.2023.102711

13. Fernández, V., Llinares-Benadero, C., Borrell, V., 2016. Cerebral cortex expansion and folding: what have we learned? EMBO J. 35, 1021–1044. 10.15252/embj.201593701

14. Fiddes, I.T., Lodewijk, G.A., Mooring, M., Bosworth, C.M., Ewing, A.D., Mantalas, G.L., Novak, A.M., Van Den Bout, A., Bishara, A., Rosenkrantz, J.L., Lorig-Roach, R., Field, A.R., Haeussler, M., Russo, L., Bhaduri, A., Nowakowski, T.J., Pollen, A.A., Dougherty, M.L., Nuttle, X., Addor, M.-C., Zwolinski, S., Katzman, S., Kriegstein, A., Eichler, E.E., Salama, S.R., Jacobs, F.M.J., Haussler, D., 2018. Human-Specific NOTCH2NL Genes Affect Notch Signaling and Cortical Neurogenesis. Cell 173, 1356–1369.e22. 10.1016/j.cell.2018.03.051

15. Fietz, S.A., Kelava, I., Vogt, J., Wilsch-Bräuninger, M., Stenzel, D., Fish, J.L., Corbeil, D., Riehn, A., Distler, W., Nitsch, R., Huttner, W.B., 2010. OSVZ progenitors of human and ferret neocortex are epithelial-like and expand by integrin signaling. Nat. Neurosci. 13, 690– 699. 10.1038/nn.2553

16. Fischer, J., Fernández Ortuño, E., Marsoner, F., Artioli, A., Peters, J., Namba, T., Eugster Oegema, C., Huttner, W.B., Ladewig, J., Heide, M., 2022. Human-specific *ARHGAP11B* ensures human-like basal progenitor levels in hominid cerebral organoids. EMBO Rep. 23, e54728. 10.15252/embr.202254728

17. Gaignerie, A., Lefort, N., Rousselle, M., Forest-Choquet, V., Flippe, L., Francois–Campion, V., Girardeau, A., Caillaud, A., Chariau, C., Francheteau, Q., Derevier, A., Chaubron, F., Knöbel, S., Gaborit, N., Si-Tayeb, K., David, L., 2018. Urine-derived cells provide a readily accessible cell type for feeder-free mRNA reprogramming. Sci. Rep. 8, 14363. 10.1038/s41598-018-32645-2

18. Gonçalves, J.C., Quintremil, S., Yi, J., Vallee, R.B., 2020. Nesprin-2 Recruitment of BicD2 to the Nuclear Envelope Controls Dynein/Kinesin-Mediated Neuronal Migration In Vivo. Curr. Biol. 30, 3116–3129.e4. 10.1016/j.cub.2020.05.091

19. Hansen, D.V., Lui, J.H., Parker, P.R.L., Kriegstein, A.R., 2010. Neurogenic radial glia in the outer subventricular zone of human neocortex. Nature 464, 554–561. 10.1038/nature08845

20. He, Z., Dony, L., Fleck, J.S., Szałata, A., Li, K.X., Slišković, I., Lin, H.-C., Santel, M., Atamian, A., Quadrato, G., Sun, J., Pașca, S.P., Human Cell Atlas Organoid Biological Network, Amin, N.D., Kelley, K.W., Bertucci, T., Temple, S., Bowles, K.R., Caporale, N., Villa, E., Testa, G., Cruceanu, C., Binder, E.B., Camp, J.G., Theis, F.J., Treutlein, B., 2024. An integrated transcriptomic cell atlas of human neural organoids. Nature 635, 690–698. 10.1038/s41586-024-08172-8

21. Herculano-Houzel, S., 2012. The remarkable, yet not extraordinary, human brain as a scaled-up primate brain and its associated cost. Proc. Natl. Acad. Sci. 109, 10661–10668. 10.1073/pnas.1201895109

22. Hu, D.J.-K., Baffet, A.D., Nayak, T., Akhmanova, A., Doye, V., Vallee, R.B., 2013. Dynein Recruitment to Nuclear Pores Activates Apical Nuclear Migration and Mitotic Entry in Brain Progenitor Cells. Cell 154, 1300–1313. 10.1016/j.cell.2013.08.024

23. Huang, W., Bhaduri, A., Velmeshev, D., Wang, S., Wang, L., Rottkamp, C.A., Alvarez-Buylla, A., Rowitch, D.H., Kriegstein, A.R., 2020. Origins and Proliferative States of Human Oligodendrocyte Precursor Cells. Cell 182, 594–608.e11. 10.1016/j.cell.2020.06.027

24. Iefremova, V., Manikakis, G., Krefft, O., Jabali, A., Weynans, K., Wilkens, R., Marsoner, F., Brändl, B., Müller, F.-J., Koch, P., Ladewig, J., 2017. An Organoid-Based Model of Cortical Development Identifies Non-Cell-Autonomous Defects in Wnt Signaling Contributing to Miller-Dieker Syndrome. Cell Rep. 19, 50–59. 10.1016/j.celrep.2017.03.047

25. Kalebic, N., Gilardi, C., Stepien, B., Wilsch-Bräuninger, M., Long, K.R., Namba, T., Florio, M., Langen, B., Lombardot, B., Shevchenko, A., Kilimann, M.W., Kawasaki, H., Wimberger, P., Huttner, W.B., 2019. Neocortical Expansion Due to Increased Proliferation of Basal Progenitors Is Linked to Changes in Their Morphology. Cell Stem Cell 24, 535–550.e9. 10.1016/j.stem.2019.02.017

26. Kunda, P., Pelling, A.E., Liu, T., Baum, B., 2008. Moesin Controls Cortical Rigidity, Cell Rounding, and Spindle Morphogenesis during Mitosis. Curr. Biol. 18, 91–101. 10.1016/j.cub.2007.12.051

27. Kyrousi, C., O’Neill, A.C., Brazovskaja, A., He, Z., Kielkowski, P., Coquand, L., Di Giaimo, R., D’ Andrea, P., Belka, A., Forero Echeverry, A., Mei, D., Lenge, M., Cruceanu, C., Buchsbaum, I.Y., Khattak, S., Fabien, G., Binder, E., Elmslie, F., Guerrini, R., Baffet, A.D., Sieber, S.A., Treutlein, B., Robertson, S.P., Cappello, S., 2021. Extracellular LGALS3BP regulates neural progenitor position and relates to human cortical complexity. Nat. Commun. 12, 6298. 10.1038/s41467-021-26447-w

28. Li, Y., Li, Z., Wang, Changliang, Yang, M., He, Z., Wang, F., Zhang, Y., Li, R., Gong, Y., Wang, B., Fan, B., Wang, Chunyue, Chen, L., Li, H., Shi, P., Wang, N., Wei, Z., Wang, Y.-L., Jin, L., Du, P., Dong, J., Jiao, J., 2023. Spatiotemporal transcriptome atlas reveals the regional specification of the developing human brain. Cell 186, 5892–5909.e22. 10.1016/j.cell.2023.11.016

29. Lindenhofer, D., Haendeler, S., Esk, C., Littleboy, J.B., Brunet Avalos, C., Naas, J., Pflug, F.G., Van De Ven, E.G.P., Reumann, D., Baffet, A.D., Von Haeseler, A., Knoblich, J.A., 2024. Cerebral organoids display dynamic clonal growth and tunable tissue replenishment. Nat. Cell Biol. 26, 710–718. 10.1038/s41556-024-01412-z

30. Liu, D.D., He, J.Q., Sinha, R., Eastman, A.E., Toland, A.M., Morri, M., Neff, N.F., Vogel, H., Uchida, N., Weissman, I.L., 2023. Purification and characterization of human neural stem and progenitor cells. Cell 186, 1179–1194.e15. 10.1016/j.cell.2023.02.017

31. Liu, H., Zhang, S.-C., 2011. Specification of neuronal and glial subtypes from human pluripotent stem cells. Cell. Mol. Life Sci. 68, 3995–4008. 10.1007/s00018-011-0770-y

32. Luxton, G.W.G., Gomes, E.R., Folker, E.S., Vintinner, E., Gundersen, G.G., 2010. Linear Arrays of Nuclear Envelope Proteins Harness Retrograde Actin Flow for Nuclear Movement. Science 329, 956–959. 10.1126/science.1189072

33. Malatesta, P., Hartfuss, E., Götz, M., 2000. Isolation of radial glial cells by fluorescent-activated cell sorting reveals a neuronal lineage. Development 127, 5253–5263. 10.1242/dev.127.24.5253

34. Martínez-Martínez, M.Á., De Juan Romero, C., Fernández, V., Cárdenas, A., Götz, M., Borrell, V., 2016. A restricted period for formation of outer subventricular zone defined by Cdh1 and Trnp1 levels. Nat. Commun. 7, 11812. 10.1038/ncomms11812

35. Matsumoto, N., Tanaka, S., Horiike, T., Shinmyo, Y., Kawasaki, H., 2020. A discrete subtype of neural progenitor crucial for cortical folding in the gyrencephalic mammalian brain. eLife 9, e54873. 10.7554/eLife.54873

36. Noctor, S.C., Flint, A.C., Weissman, T.A., Dammerman, R.S., Kriegstein, A.R., 2001. Neurons derived from radial glial cells establish radial units in neocortex. Nature 409, 714– 720. 10.1038/35055553

37. Nowakowski, T.J., Pollen, A.A., Sandoval-Espinosa, C., Kriegstein, A.R., 2016. Transformation of the Radial Glia Scaffold Demarcates Two Stages of Human Cerebral Cortex Development. Neuron 91, 1219–1227. 10.1016/j.neuron.2016.09.005

38. Ostrem, B.E.L., Lui, J.H., Gertz, C.C., Kriegstein, A.R., 2014. Control of Outer Radial Glial Stem Cell Mitosis in the Human Brain. Cell Rep. 8, 656–664. 10.1016/j.celrep.2014.06.058

39. Pilaz, L.-J., Patti, D., Marcy, G., Ollier, E., Pfister, S., Douglas, R.J., Betizeau, M., Gautier, E., Cortay, V., Doerflinger, N., Kennedy, H., Dehay, C., 2009. Forced G1-phase reduction alters mode of division, neuron number, and laminar phenotype in the cerebral cortex. Proc. Natl. Acad. Sci. 106, 21924–21929. 10.1073/pnas.0909894106

40. Pinson, A., Xing, L., Namba, T., Kalebic, N., Peters, J., Oegema, C.E., Traikov, S., Reppe, K., Riesenberg, S., Maricic, T., Derihaci, R., Wimberger, P., Pääbo, S., Huttner, W.B., 2022. Human TKTL1 implies greater neurogenesis in frontal neocortex of modern humans than Neanderthals. Science 377, eabl6422. 10.1126/science.abl6422

41. Qian, X., Coleman, K., Jiang, S., Kriz, A.J., Marciano, J.H., Luo, C., Cai, C., Manam, M.D., Caglayan, E., Otani, A., Ghosh, U., Shao, D.D., Andersen, R.E., Neil, J.E., Johnson, R., LeFevre, A., Hecht, J.L., Miller, M.B., Sun, L., Stringer, C., Li, M., Walsh, C.A., 2024. Spatial Single-cell Analysis Decodes Cortical Layer and Area Specification. 10.1101/2024.06.05.597673

42. Qian, X., Nguyen, H.N., Song, M.M., Hadiono, C., Ogden, S.C., Hammack, C., Yao, B., Hamersky, G.R., Jacob, F., Zhong, C., Yoon, K., Jeang, W., Lin, L., Li, Y., Thakor, J., Berg, D.A., Zhang, C., Kang, E., Chickering, M., Nauen, D., Ho, C.-Y., Wen, Z., Christian, K.M., Shi, P.-Y., Maher, B.J., Wu, H., Jin, P., Tang, H., Song, H., Ming, G., 2016. Brain-Region-Specific Organoids Using Mini-bioreactors for Modeling ZIKV Exposure. Cell 165, 1238– 1254. 10.1016/j.cell.2016.04.032

43. Qian, X., Su, Y., Adam, C.D., Deutschmann, A.U., Pather, S.R., Goldberg, E.M., Su, K., Li, S., Lu, L., Jacob, F., Nguyen, P.T.T., Huh, S., Hoke, A., Swinford-Jackson, S.E., Wen, Z., Gu, X., Pierce, R.C., Wu, H., Briand, L.A., Chen, H.I., Wolf, J.A., Song, H., Ming, G., 2020. Sliced Human Cortical Organoids for Modeling Distinct Cortical Layer Formation. Cell Stem Cell 26, 766–781.e9. 10.1016/j.stem.2020.02.002

44. Ramanathan, S.P., Helenius, J., Stewart, M.P., Cattin, C.J., Hyman, A.A., Muller, D.J., 2015. Cdk1-dependent mitotic enrichment of cortical myosin II promotes cell rounding against confinement. Nat. Cell Biol. 17, 148–159. 10.1038/ncb3098

45. Reillo, I., De Juan Romero, C., García-Cabezas, M.Á., Borrell, V., 2011. A Role for Intermediate Radial Glia in the Tangential Expansion of the Mammalian Cerebral Cortex. Cereb. Cortex 21, 1674–1694. 10.1093/cercor/bhq238

46. Reiner, O., Carrozzo, R., Shen, Y., Wehnert, M., Faustinella, F., Dobyns, W.B., Caskey, C.T., Ledbetter, D.H., 1993. Isolation of a Miller–Dicker lissencephaly gene containing G protein β-subunit-like repeats. Nature 364, 717–721. 10.1038/364717a0

47. Romero, D.M., Bahi-Buisson, N., Francis, F., 2018. Genetics and mechanisms leading to human cortical malformations. Semin. Cell Dev. Biol. 76, 33–75. 10.1016/j.semcdb.2017.09.031

48. Sauerland, C., Menzies, B.R., Glatzle, M., Seeger, J., Renfree, M.B., Fietz, S.A., 2018. The Basal Radial Glia Occurs in Marsupials and Underlies the Evolution of an Expanded Neocortex in Therian Mammals. Cereb. Cortex 28, 145–157. 10.1093/cercor/bhw360

49. Serres, M.P., Samwer, M., Truong Quang, B.A., Lavoie, G., Perera, U., Görlich, D., Charras, G., Petronczki, M., Roux, P.P., Paluch, E.K., 2020. F-Actin Interactome Reveals Vimentin as a Key Regulator of Actin Organization and Cell Mechanics in Mitosis. Dev. Cell 52, 210–222.e7. 10.1016/j.devcel.2019.12.011

50. Sloan, S.A., Andersen, J., Pașca, A.M., Birey, F., Pașca, S.P., 2018. Generation and assembly of human brain region-specific three-dimensional cultures. Nat. Protoc. 13, 2062–2085. 10.1038/s41596-018-0032-7

51. Suzuki, I.K., Gacquer, D., Van Heurck, R., Kumar, D., Wojno, M., Bilheu, A., Herpoel, A., Lambert, N., Cheron, J., Polleux, F., Detours, V., Vanderhaeghen, P., 2018. Human-Specific NOTCH2NL Genes Expand Cortical Neurogenesis through Delta/Notch Regulation. Cell 173, 1370–1384.e16. 10.1016/j.cell.2018.03.067

52. Taubenberger, A.V., Baum, B., Matthews, H.K., 2020. The Mechanics of Mitotic Cell Rounding. Front. Cell Dev. Biol. 8, 687. 10.3389/fcell.2020.00687

53. Toyoda, Y., Cattin, C.J., Stewart, M.P., Poser, I., Theis, M., Kurzchalia, T.V., Buchholz, F., Hyman, A.A., Müller, D.J., 2017. Genome-scale single-cell mechanical phenotyping reveals disease-related genes involved in mitotic rounding. Nat. Commun. 8, 1266. 10.1038/s41467-017-01147-6

54. Tsai, J.-W., Bremner, K.H., Vallee, R.B., 2007. Dual subcellular roles for LIS1 and dynein in radial neuronal migration in live brain tissue. Nat. Neurosci. 10, 970–979. 10.1038/nn1934

55. Tsai, J.-W., Lian, W.-N., Kemal, S., Kriegstein, A.R., Vallee, R.B., 2010. Kinesin 3 and cytoplasmic dynein mediate interkinetic nuclear migration in neural stem cells. Nat. Neurosci. 13, 1463–1471. 10.1038/nn.2665

56. Uzquiano, A., Kedaigle, A.J., Pigoni, M., Paulsen, B., Adiconis, X., Kim, K., Faits, T., Nagaraja, S., Antón-Bolaños, N., Gerhardinger, C., Tucewicz, A., Murray, E., Jin, X., Buenrostro, J., Chen, F., Velasco, S., Regev, A., Levin, J.Z., Arlotta, P., 2022. Proper acquisition of cell class identity in organoids allows definition of fate specification programs of the human cerebral cortex. Cell 185, 3770–3788.e27. 10.1016/j.cell.2022.09.010

57. Walsh, R.M., Luongo, R., Giacomelli, E., Ciceri, G., Rittenhouse, C., Verrillo, A., Galimberti, M., Bocchi, V.D., Wu, Y., Xu, N., Mosole, S., Muller, J., Vezzoli, E., Jungverdorben, J., Zhou, T., Barker, R.A., Cattaneo, E., Studer, L., Baggiolini, A., 2024. Generation of human cerebral organoids with a structured outer subventricular zone. Cell Rep. 43, 114031. 10.1016/j.celrep.2024.114031

58. Wang, L., Wang, C., Moriano, J.A., Chen, S., Zuo, G., Cebrián-Silla, A., Zhang, S., Mukhtar, T., Wang, S., Song, M., De Oliveira, L.G., Bi, Q., Augustin, J.J., Ge, X., Paredes, M.F., Huang, E.J., Alvarez-Buylla, A., Duan, X., Li, J., Kriegstein, A.R., 2024. Molecular and cellular dynamics of the developing human neocortex at single-cell resolution. 10.1101/2024.01.16.575956

59. Wang, R., Sharma, R., Shen, X., Laughney, A.M., Funato, K., Clark, P.J., Shpokayte, M., Morgenstern, P., Navare, M., Xu, Y., Harbi, S., Masilionis, I., Nanjangud, G., Yang, Y., Duran-Rehbein, G., Hemberg, M., Pe’er, D., Tabar, V., 2020. Adult Human Glioblastomas Harbor Radial Glia-like Cells. Stem Cell Rep. 14, 338–350. 10.1016/j.stemcr.2020.01.007

60. Wimmer, R., Baffet, A.D., 2023. The microtubule cytoskeleton of radial glial progenitor cells. Curr. Opin. Neurobiol. 80, 102709. 10.1016/j.conb.2023.102709

61. Xie, Y., Bergström, T., Jiang, Y., Johansson, P., Marinescu, V.D., Lindberg, N., Segerman, A., Wicher, G., Niklasson, M., Baskaran, S., Sreedharan, S., Everlien, I., Kastemar, M., Hermansson, A., Elfineh, L., Libard, S., Holland, E.C., Hesselager, G., Alafuzoff, I., Westermark, B., Nelander, S., Forsberg-Nilsson, K., Uhrbom, L., 2015. The Human Glioblastoma Cell Culture Resource: Validated Cell Models Representing All Molecular Subtypes. EBioMedicine 2, 1351–1363. 10.1016/j.ebiom.2015.08.026

62. Xing, L., Gkini, V., Nieminen, A.I., Zhou, H.-C., Aquilino, M., Naumann, R., Reppe, K., Tanaka, K., Carmeliet, P., Heikinheimo, O., Pääbo, S., Huttner, W.B., Namba, T., 2024. Functional synergy of a human-specific and an ape-specific metabolic regulator in human neocortex development. Nat. Commun. 15, 3468. 10.1038/s41467-024-47437-8

63. Zillich, L., Rossetti, A.C., Fechtner, O., Gasparotto, M., Maillard, C., Hoffrichter, A., Zillich, E., Jabali, A., Marsoner, F., Wilkens, R., Schroeter, C.B., Hentschel, A., Meuth, S.G., Ruck, T., Koch, P., Roos, A., Bahi-Buisson, N., Francis, F., Ladewig, J., 2022. Unraveling LIS1-Lissencephaly: Insights from Cerebral Organoids Suggest Severity-Dependent Genotype-Phenotype Correlations, Molecular Mechanisms and Therapeutic Strategies. 10.1101/2022.12.19.520907

